# Differential regulation of the proteome and phosphosproteome along the dorso-ventral axis of the early *Drosophila* embryo

**DOI:** 10.1101/2023.08.24.554590

**Authors:** Juan Manuel Gomez, Hendrik Nolte, Elisabeth Vogelsang, Bipasha Dey, Michiko Takeda, Girolamo Giudice, Miriam Faxel, Alina Cepraga, Robert Patrick Zinzen, Marcus Krüger, Evangelia Petsalaki, Yu-Chiun Wang, Maria Leptin

## Abstract

The initially homogeneous epithelium of the early *Drosophila* embryo differentiates into regional subpopulations with different behaviours and physical properties that are needed for morphogenesis. The factors at top of the genetic hierarchy that control these behaviours are known, but many of their targets are not. To understand how proteins work together to mediate differential cellular activities, we studied in an unbiased manner the proteomes and phosphoproteomes of the three main cell populations along the dorso-ventral axis during gastrulation using mutant embryos that represent the different populations. We detected 6111 protein groups and 6259 phosphosites of which 3399 and 3433 respectively, were differentially regulated. The changes in phosphosite abundance did not correlate with changes in host protein abundance, showing phosphorylation to be a regulatory step during gastrulation. Hierarchical clustering of protein groups and phosphosites identified clusters that contain known fate determinants such as Doc1, Sog, Snail and Twist. The recovery of the appropriate known marker proteins in each of the different mutants we used validated the approach, but also revealed that two mutations that both interfere with the dorsal fate pathway, *Toll^10B^* and *serpin27a^ex^* do this in very different manners. Diffused network analyses within each cluster point to microtubule components as one of the main groups of regulated proteins. Functional studies on the role of microtubules provide the proof of principle that microtubules have different functions in different domains along the DV axis of the embryo.

## Introduction

Morphogenesis is the developmental process that creates the three-dimensional morphology of tissues. The first morphogenetic event in metazoans is gastrulation, in which an epithelium gives rise to the germ layers from which all adult tissues derive. *Drosophila* gastrulation is probably one of the best studied embryo-scale morphogenetic processes: it is initiated by the formation of a ventral furrow that leads to the internalisation of the mesoderm. The internalisation of the mesoderm causes the ventral displacement of the neuroectoderm, the ectodermal cell population on the lateral side of the embryo, in the absence of particular cell shape behaviours. Finally, this ventral displacement of the neuroectoderm is accommodated by the stretching of dorsal ectodermal cells [1, 2]. Therefore, the behaviour of these cell populations can be used to study the connection between cell fate and cell shape regulation.

The behaviour of a cell is determined by the identity and the state of the proteins within the cell, and by the networks through which these proteins interact. The first step to fill the gap between cell fate and cell shape behaviour is to understand how the embryonic cell populations differ in their biochemical composition. Most of the cellular components pre-exist in the egg, having been provided maternally during oogenesis either as RNA or as protein. With the exception of the determinants for anterior-posterior (AP) and dorso-ventral (DV) patterning most of these proteins are distributed throughout the early embryo. As differentiation proceeds, they may be acted upon in a region-specific manner [3]. For example, adherens junctions and the acto-myosin meshwork are dramatically remodelled in ventral cells [2, 4].

The mechanism by which the differentiation of embryonic cell populations is controlled is understood in great depth, largely through the study of mutants. Briefly, a gradient of the transcription factor Dorsal with its high point in nuclei on the ventral side is triggered by a graded extracellular signal that is transmitted through the transmembrane receptor Toll [5, 6]. We use for our work here mutations in three genes that control dorso-ventral fates, *Toll, serpin27A* and *gastrulation defective.* Female flies that are homozygous for certain alleles of these mutations, or combinations of alleles, lay eggs that develop into embryos in which all cells express genes characteristic for only one domain of the normal embryo - either the ventral domain, or the lateral or the dorsal domain, and to which we refer here as ventralized, lateralized or dorsalized.

The transcription factors and signalling cascades set up by DV patterning and their downstream target proteins then act upon some of the maternally provided proteins in a region-specific manner. Among protein-level post-translational modifications, phosphorylation is fast and reversible and plays key roles during early embryogenesis: from regulating elements in the Toll and Dpp pathways, to the activation of the Rho Pathway within the mesoderm [5, 7]. Therefore, phosphorylation is likely to be at least one way of also regulating cell behaviours along the dorso-ventral axis in a cost-effective and timely manner.

Differences between embryonic cell populations along the DV axis have been studied with transcriptomic and proteomic methods [8–11] but with limited depth and temporal resolution. Studies looking at changes over time identified proteins that appear during the maternal to zygotic transition [12, 13] and later in embryogenesis [14, 15], but had no spatial or cell type specificity. None of these studies addressed the region-specific post-translational regulation of proteins.

To identify missing links in the pathways from known cell fate determining factors and region-specific cell behaviours, we analysed the proteomes and the phosphoproteomes of mutants representing different cell populations along the dorso-ventral axis of the embryo. We find many proteins with differences in abundance across the populations that do not show the same differences in RNA abundance. We also find region-specific phosphorylation patterns in proteins that are ubiquitously expressed. Networks of phosphoproteins enriched in specific populations included proteasome components, RNA stress granules/P-bodies, adherens junctions associated proteins and microtubule components/associated proteins. A proof of principle test of the role of microtubules in the gastrulating embryos and revealed differential functions in the cell populations along the DV axis.

## Results

### 1) Biological validation of dorso-ventral patterning mutants as representatives of dorso-entral cell populations in the wild type embryo

To study the proteomes and the phosphoproteomes of cell populations in the early embryo (Supplementary Fig. 1A), we used mutants in which all cells in the embryo represent only one subset of the cell types present in the wild type embryo. Because we were interested in cell behaviours that affect the first step of gastrulation, which is driven by differences in cell behaviour along the DV axis, we used mutants for genes of the DV patterning pathway. The embryos were derived from mothers mutant for the genes *gastrulation defective* (gd), *Toll* (Tl), or *Serpin27A* (spn27A). These embryos represent dorsal, lateral and ventral cell populations as judged by the expression patterns of D-V fate determining genes (Fig 1A, Fig 1A, Supplementary Table 1, Supplementary Fig. 1B). ‘Lateralized’ and ‘dorsalized’ embryos from *Tl^rm10^/Tl^rm9^* and *gd^9^*/*gd^9^* mothers express neither *twist* nor *snail*, whereas ventralized embryos from *Toll^10B^/def* and *spn27A^ex^/def* mothers express *twist* and *snail* around their entire circumference in the trunk region (Fig. 1B and Supplementary Table 1). In embryos from *Tl^rm10^/Tl^rm9^* mothers, Sog expression expands dorsally and ventrally, whereas Dpp expression expands ventrally (Fig. 1B and Supplementary Table 1). These expression patterns show some variation and are not entirely homogeneous: ventralized embryos often have a gap in *snail* expression in a small dorsal-anterior domain around the procephalic furrow. In this region, we detected *sog* expression instead, suggesting ventralized embryos retain a neuroectodermal fate in a restricted area of the embryo (Fig. 1B, *sog* probe).

**Figure 1.**
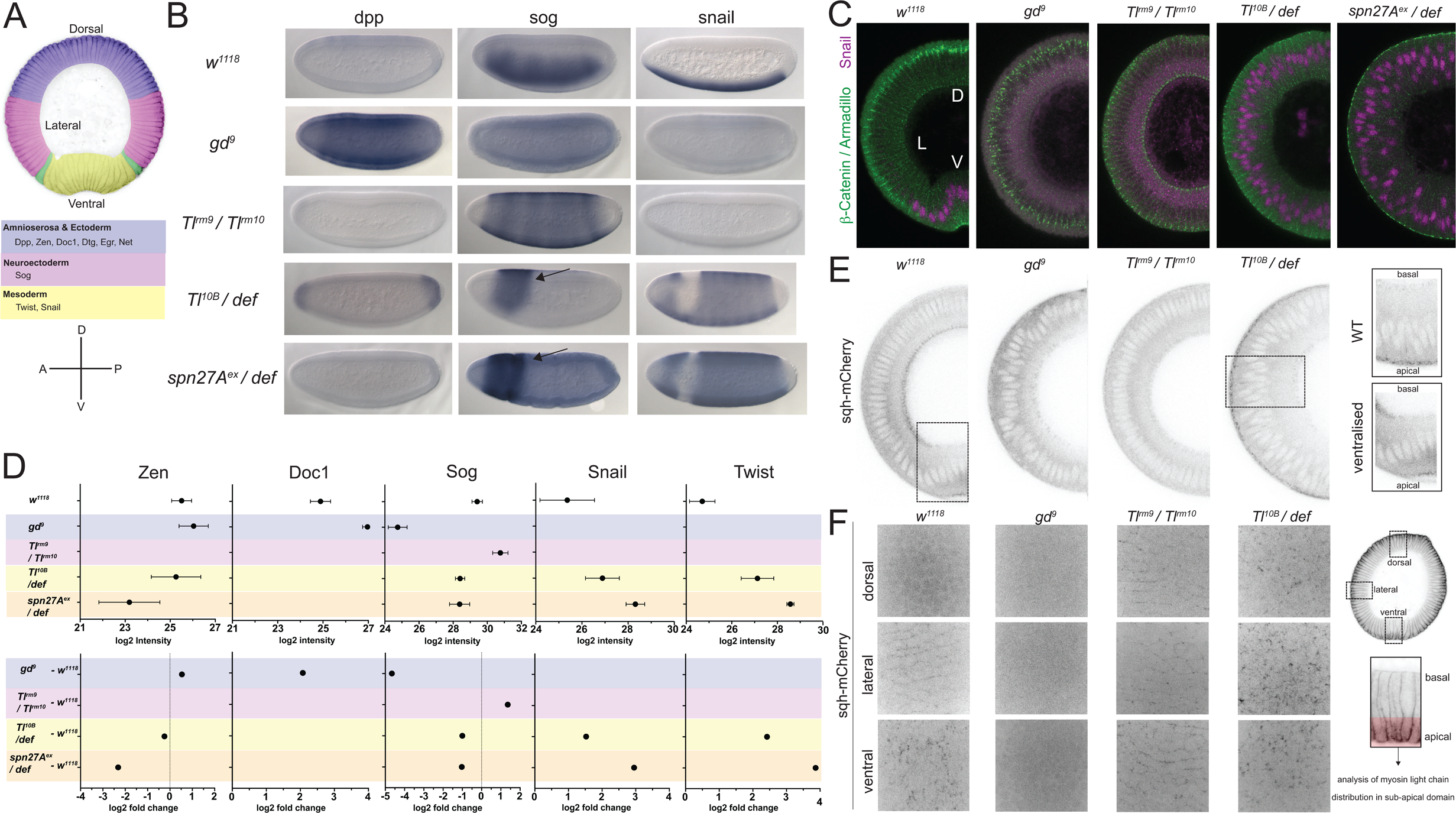
Validation of mutants as representatives of embryonic cell populations. **A)** Top: colour-coded schematic of the cell populations along the dorso-ventral axis of a *Drosophila* embryo during gastrulation: blue, ‘dorsal’: ectoderm and amnioserosa; magenta, ‘lateral’: neuroectoderm; green, mesectoderm; yellow ‘ventral’: mesoderm. Middle: examples of dorso-ventral fate determinants of the domains shown in top panel. Bottom: dorso-ventral and anterior-posterior axes for reference in panel B. **B)** RNA *in situ* hybridisations using probes for genes expressed in dorsal (*dpp*), lateral (*sog*) and ventral (*snail*) cell populations in wild type (*w^1118^*) and embryos derived from mothers mutant for dorso-ventral patterning genes (*gd^9^,* dorsalized, *Tl^rm9^*/ *Tl^rm10^* lateralized, *Toll^10B^*/*def* and *spn27A^ex^*/*def* both ventralized). Notice the expansion of *dpp, sog* and *snail* expression in dorsalized, lateralized and ventralized embryos respectively. Arrow indicates reminiscent neuroectodermal polarity in ventralized embryos. **C)** Images (confocal, max-projected) of physical cross-sections from heat-fixed embryos stained using antibodies against β-Catenin/Armadillo (green) and Snail (magenta). D is dorsal domain; L is lateral domain; V is ventral domain). **D)** Log2 intensity (top) and log2 fold change (FC, bottom) of proteins in wild type, dorsalized (blue), lateralized (magenta) and ventralized (yellow) embryos. Bars depict mean and standard error of the mean across replicates. Absence of a dot indicates the protein was not detected or Log2FC calculation not feasible; absence of error bars in log2 intensity indicates protein was detected only in a single biological replicate. Dotted line indicates Log2FC = 0. **E)** Cross-section images (two-photon, single sections) showing Myosin Light Chain (sqh-mCherry) distribution in living wild type, dorsalized, lateralized and ventralized embryos. Insets show magnified ectopic sqh-mCherry signal distribution in wild type vs. ventralized embryos. **F)** Images (spinning disk, max-projected) showing myosin distribution in the sub-apical domain of living wild type, dorsalized, lateralized and ventralized embryos along their dorso-ventral axis.

Because we want to use these mutants to identify proteins that reflect or control differential cell behaviour it was important to ascertain that the cells in these mutants recapitulate faithfully the biological qualities of the corresponding cell populations in the wild type embryo [2, 4], specifically of the localisation of the adherens junctions and the cortical actomyosin meshwork. We find that, as in the mesoderm of wildtype embryos, the adherens junctions (as visualised by immunostaining for Armadillo/μ-Catenin; Fig. 1C) relocalize apically in the ventralized mutants, but remain apico-lateral in lateralized and shift slightly more basally in dorsalized mutants, again mirroring the morphology of lateral and dorsal regions of the wildtype embryo (Supplementary Fig. 1A, Fig. 1C). Similarly, the apical actomyosin network, which we characterised in living embryos expressing a fluorescently tagged myosin light chain (sqh-mCherry, Fig. 1E,F) forms a pulsatile apical network in ventralized embryos, whereas myosin accumulates at cell junctions in lateralized embryos, and dorsalized embryos dissolve the loose apical actomyosin of the early blastoderm (Fig. 1E,F).

In summary, in terms of marker gene expression and cell behaviour, the cells in these mutants resemble the corresponding embryonic cell populations of a wild type embryo, showing that these mutant cell populations are good sources of material to analyse the proteomic and phosphoproteomic composition of the natural cell populations at the onset of gastrulation.

### 2 ) The proteome and the phosphoproteome of four cell populations during gastrulation

To study the proteins and the phosphosites that might be relevant for cell behaviour during gastrulation, we focused on a narrow developmental time window for sample collection. We synchronised egg collections and manually collected embryos from wild type and mutant mothers aged for 165-180 minutes after egg deposition at 25°C (Stage 6, Supplementary Fig. 1A,B). We analysed their peptides and phospho-peptides with unbiased label-free quantification (LFQ) and SILAC (Stable Isotope Labelling with Amino acids in Culture [16–18]. Wild type larvae were raised with SILAC yeast as the only source of amino acids, resulting in ∼75% of labelled proteome in the progeny of the emerging female flies (Supplementary Fig. 1C, D). SILAC labelling did not affect the phosphoproteome of wild type embryos, and had only a minor effect on phosphosite intensity distribution, indicating standardisation with Lys 13/6 was a valid approach (Supplementary Fig. 1E).

In the proteomic analyses, we identified 6111 protein groups (to which, for the sake of simplicity, we will refer simply as proteins) across all genotypes. 5883 of these were detected in wild type embryos (Fig 2A), exceeding the previously reported number of protein groups found in early *Drosophila* embryogenesis (0-2hs: 4825; 2-4hs: 5064 proteins [15]. A small number (519/6111) were detected in individual or multiple genotypes, but not in all (Fig. 2B). This category included the DV fate determinants Doc1, Snail, Twist and dMyc (Fig. 1A,B). The phosphoproteomic analysis identified 6259 phosphosites distributed over 1847 proteins (Fig. 2C). The percentage of phosphosites detected in subsets but not all genotypes (30%, Fig. 2D) was larger than for the proteome, even though the number of proteins and the number of phosphosites was similar across all genotypes (Fig. 2A,C). 28% of the proteins (1699/6111) and 9% of the phosphosites (573/6259) differed significantly in an ANOVA test across all five populations (wild type and four mutants) (permutation-based FDR < 0.1, s0=0.1).

**Figure 2.**
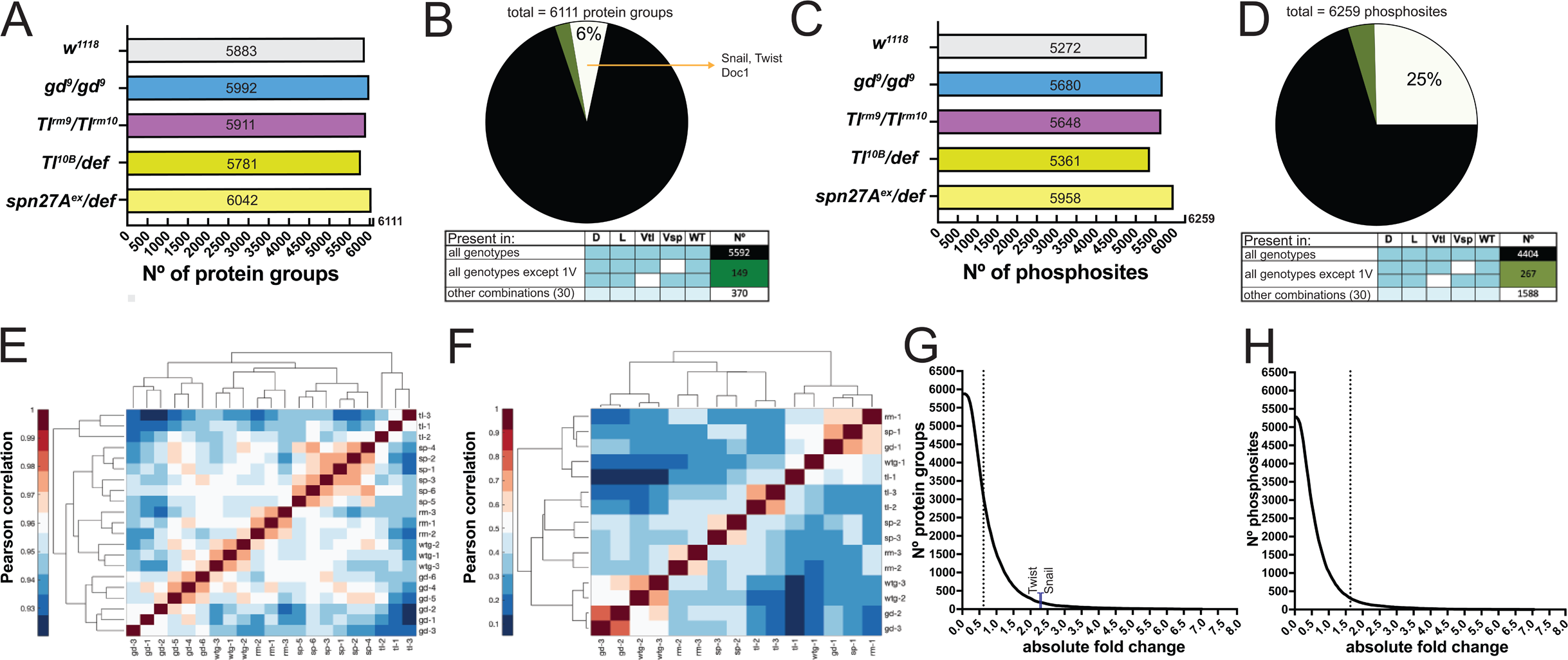
Proteomes and phosphoproteomes of wildtype and mutant embryos. **A and C)** Number of protein groups (A) or phosphosites (C) detected in wildtype, dorsalized (*gd^9^*), lateralized (*Tl^rm9^*/ *Tl^rm10^*), and ventralized embryos (*Toll10B*/*def* and *spn27A^ex^*/*def*). **B and D)** Intersection analysis of detected protein groups (B) or phosphosites (D). Black: detected in at least 1 replicate in all genotypes; green: detected in at least 1 replicate in all genotypes except 1 ventralized condition; white: detected in at least 1 replicate in any other combination. **E and F)** Correlation matrix between the replicates of the proteomic (E) and phosphoproteomic (F) experiments using the Pearson correlation coefficient. Protein groups and phosphosites detected in all of the replicates in all of the genotypes were used to construct the correlation matrices. **G and H)** Distribution of the number of protein groups (G) or phosphosites (H) exceeding an absolute fold change (vs. wild type, in log2 scale). Dotted line depicts the absolute fold change corresponding to 50% of the analysed protein groups (G) or phosphosites (H).

We determined the degree of experimental variability by generating correlation matrices both for the proteome and the phosphoproteome. For the proteome, the replicates from the same genotypes clustered together (Fig. 2E). For the phosphoproteome, the first replicate of each genotype was separated from the other two replicates (Fig. 2F). We nevertheless kept all replicates for further analyses because it was impossible to determine the experimental source for this variation.

The enrichment for proteins or phosphosites in the mutant genotypes over the wild-type ranged from near-zero to 100 fold (Fig. 2G,H). Half of the proteins or phosphosites changed by less than 1.5-fold. The fold-changes for the mesodermal fate determinants Snail (5.4 fold) and Twist (14.6 fold) were the largest among the DV fate determinants (Fig. 2G); fifty-five of the entire set of proteins showed a stronger change versus wildtype than Twist.

To test if the recovered protein populations represented the cell populations in the embryo, we analysed whether they contained known marker proteins. We first looked for the protein product of the gene that was mutated in each group of embryos. We detected both Toll and Spn27A. In *Tl^rm9^/Tl^rm10^* embryos, the Tl protein was decreased in abundance and Spn27A was less abundant in embryos derived from *spn27A^ex^/df* mothers (Supplementary Fig. 2A).

Proteins that are known to be expressed differentially along the DV axis (Fig. 1A) were found with increased abundance in the appropriate genotypes: Snail,Twist, Mdr49, Traf4 and CG4500 in ventralized embryos; pro-neuroectodermal (lateral) factor Sog in lateralized embryos; pro-ectodermal (dorsal) factors, such as Zen, Doc1, Dtg, Net and Egr in dorsalized embryos (Fig. 1D, Supplementary Fig. 2C, D). We also identified phosphosites known to be regulated along the DV axis during gastrulation. This included phosphosites in Toll and three serines phosphorylated by CKII in positions 463, 467 and 468 [19] (Supplementary Fig. 2E). Toll was differentially phosphorylated in lateralized and ventralized embryos in serine-871 (Supplementary Fig. 2B), consistent with reported Tl phosphorylation by Pelle [20]. Phosphosites in proteins associated with the Rho pathway will be discussed below. In summary, the proteomic and phosphoproteomic screen correctly identified known differentially expressed proteins and phosphosites.

### 3) A linear model for quantitative interpretation of the proteomes

Having established that each population contained the correct marker proteins, we could now ask which unknown proteins were changed. Our knowledge of the genetics of the dorso-ventral patterning system gives us additional criteria that we can use to analyse the data in a stringent manner. We know that the changes in the protein sets typical for the three populations along the DV axis should change in concert in a well-controlled manner in all of the mutants. Rather than simply looking for individual pair-wise changes, we can, and must, therefore impose this as an additional criterion in determining any potential proteins of interest: each protein must change in a manner that ‘makes sense’ genetically.

The assumption that each mutant represents a defined region of the embryo makes a simple prediction for the expected outcome of the measurements: the amounts of protein found in each of the mutants, normalised to the area of the embryo the corresponding region occupies, should add up to the amount of protein in the wildtype. For example, the transcription factor Snail is expressed only in the prospective mesoderm (ventral domain) in the wildtype embryo, but along the entire DV axis of ventralized embryos, and it is absent in lateralized and dorsalized embryos (Fig. 1B, Supplementary Fig. 3A). This is also reflected correctly in the proteomes: Snail is absent in the dorsalized and lateralized proteomes, and its level is higher in the proteomes from the ventralized embryos. Thus, Snail shows an ideal behaviour in each of the DV mutant genotypes because it recapitulates the expression of Snail in the corresponding domains of a wild type embryo.

We developed a ‘linear model’ that is based on this additional genetic criterion as a means to evaluate simultaneously all DV mutant genotypes. We calculated for each protein whether it followed the rule that its level in the mutants should add up to the wildtype level by comparing that sum to its real abundance measured in the wild type embryo. The mesoderm represents approximately 20% of the circumference of the embryo [21] the lateral ectoderm, as defined by the lateral expression domains of genes such as sog, rhomboid or brinker covers 20% of the circumference on each side [11, 22], which leaves 40% for the dorsal side.

To obtain the theoretical abundance (^t^wt) for each protein we added the mean intensity measurements (normalised, see methods) for each mutant weighted by these values (0.2 for the mesoderm, 0.4 for the lateral ectoderm, 0.4 for the dorsal region):

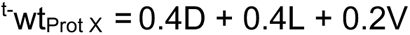

We determined the deviation of the theoretical abundance (^t^wt) from the measured abundance for the wildtype (^m^wt) as log2 of their ratio:

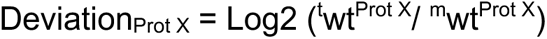

A Log2 value of 0 for the deviation therefore indicates a perfect match between the theoretical calculation and the actual measurement. When we apply this analysis to one of the marker proteins, Snail, we arrive at a deviation value of 0.1 (1.07 in linear scale) in the case of the data set for *Toll^10B^*. This shows that for this protein, the mutants represent the regional distribution in the wildtype very well. With the dataset for *spn27A*, the deviation value for Snail is 1.55 (2.91 in linear scale), which indicates that this mutant genotype over-represents the ventral population, if used with the assumed weightings for the populations

We tested what ratio of the areas of the embryonic regions would give the best fit of the theoretical sum with the data in the wildtype without any prior assumption about their real area in the embryo. A parameter screen of the entire range of possible values (0 to 1 for each mutant population) yielded good fits for a range of combinations. For the two best, the area for the ventral region was larger than observed *in vivo*, while the third best was one for which the ventral domain corresponded to the experimentally measured value of 0.2 as did the lateral and dorsal values (Supplementary Table 4, Supplementary Fig. 3B,C)]. We therefore chose this set to calculate the values for all proteins.

We found that the majority of the proteins had a deviation around zero (Fig. 3A). This would in fact be expected for any proteins that are expressed ubiquitously in the wild type and therefore present in equal amounts in all genotypes. Indeed most proteins in the early embryo are deposited by the mother during the production of the egg and are therefore present globally. Their calculated weighted sum would always be the same as the wildtype value. By contrast, this would not necessarily be expected for the proteins that show significant differences between at least two mutant conditions, i.e. the statistically significant subset. Yet, most of these also fall into the range between −0.5 and +0.5, i.e. less than 1.4 fold deviation (Fig. 3A, Supplementary Table 5). This shows that the majority of proteins fit the linear model, which in turn indicates that the mutant values are good representations of protein abundance in the corresponding domains of a wild type embryo. The proteins with the most extreme deviations (more than two-fold) did not come from any well-defined class of proteins, but represented a wide range of protein ontologies (Supplementary Fig. 3D, Supplementary Table 12).

**Figure 3.**
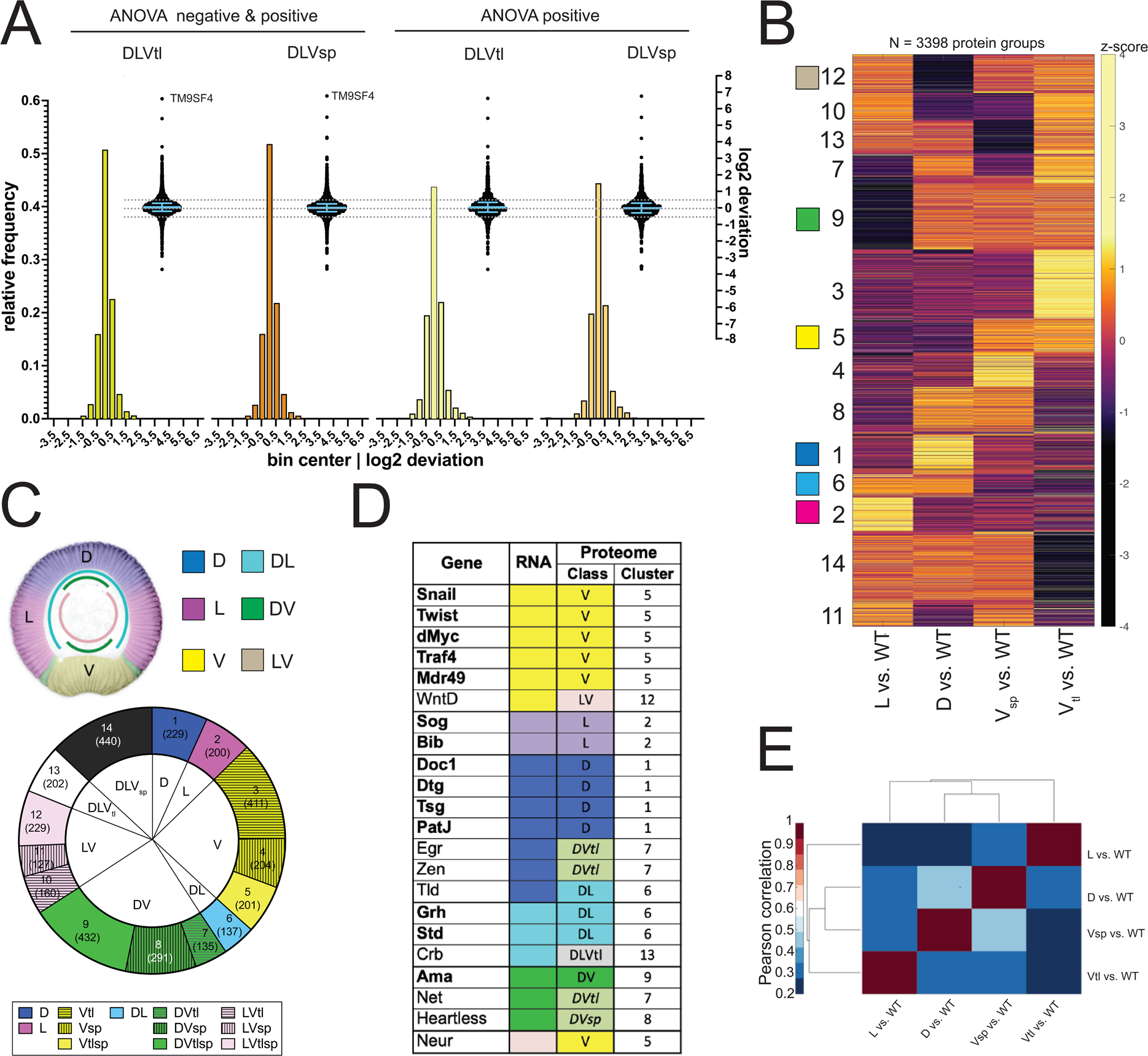
Analysis of the proteomes. **A)** Two different representations, a histogram and a swarm plot, of the deviation parameter (in log2 scale) calculated for each of the two ventralized genotypes (D: dorsalized, L: lateralized, Vtl: *Toll^10B^*/*def*, Vsp: *spn27A^ex^*/*def*). This was done once for all proteins present in all genotypes, and once only for those that were ANOVA positive. In the swarm plot, each dot represents a protein. The y-axis for the histograms is shown on the left, for swarm plots on the right. Blue bars show the median with inter-quartile range (IQR). The median is close to zero and the IQRs range from −0.18 to 0.3. The dotted line indicates the 0.5 and −0.5 deviation in swarn plots. Histograms and swarm plots were assembled with proteins detected in all genotypes. **B)** Clustergram of the hierarchical clustering (dendrograms not shown) of 3398 filtered proteins. Z-scores were calculated using the thresholded fold changes between DV mutants and wild type. Numbers identify the different clusters for reference across panels and figures. Coloured boxes indicate the DV clusters that show consistent behaviour for the two ventralising genotypes (for colour-coding see panel C). **C)** Top: Regulation categories emerging from hierarchical clustering: D (blue), increased abundance in dorsal domain; L (magenta), increased abundance in lateral domain; V (yellow), increased abundance in ventral domain; DL (cyan) increased abundance in dorsal and lateral domains; DV (green), increased abundance in dorsal and ventral domains; LV (pink) increased abundance in lateral and ventral domains. Bottom: Pie chart showing the number of protein groups (in brackets) allocated to each cluster in (B), grouped by their regulation category. **D)** Genes with restricted dorso-ventral expression with their reported RNA expression pattern, their allocation to proteome clusters (numbers refer to clusters in panel B and C), and their regulation category along the dorso-ventral axis. **E)** Correlation matrices using Pearson correlation coefficient between the fold changes of each DV mutant vs. wild type in the proteome experiment.

### 4) Hierarchical clustering strategy and emerging regulation categories from the integrated proteomes

To find the proteins that function in a tissue-specific manner during gastrulation we sorted the proteins into sets that change in concert in all of the mutants in the predicted, ‘correct’ manner, again using the assumptions that underlie this study, i.e. that the changes in the different mutants would be expected to correlate with each other in logical ways, as described above.

Rather than focusing only on the proteins that the ANOVA had shown as significantly modulated, we included in this analysis all proteins that were detectable in the wildtype (5883/6111), even if they were undetectable in one or more mutant populations. This allows us to include the important group of proteins that show a ‘perfect’ behaviour, like Twist, Snail or Doc1, in that they are undetectable in the mutants that correspond to the regions in the normal embryo where these genes are not expressed.

We used hierarchical clustering to identify the sets of proteins that change in the mutants in the same manner. For this analysis, we ignored the quantitative extent of the changes in the mutants versus the wildtype, and only focused on the direction of change. Thus, we assigned a value of −1 to any negative change against wildtype, +1 to any positive change, and 0 for ‘no change’ with a +/-1.4 fold change as a threshold for ‘no change’. ‘Undetectable’ in a mutant implies ‘reduced vs. wildtype’ and therefore scored as −1. For example, Snail is undetectable in D and L and increased in V, and therefore has the values D = −1, L = −1 and V = 1. We did not include in our further steps those proteins that were unchanged in all genotypes (2156/6111) or those that were either upregulated or downregulated versus the wildtype in all mutants (329/6111).

We clustered the remaining 3398 proteins. Instead of the raw assignments of −1, 0 and +1, we used the Z-scores for each protein. The reason for this is that this method takes into account that value sets that represent similar relative differences between the mutants, for example, 0 −1 −1 vs. 1 −1 −1 or 1, 0, 0 are biologically more similar to each other than the raw values indicate. The z-scores for all of these cases would be 1.1547 −0.5774 −0.5774.

Based on known gene expression patterns across the DV axis one would expect 6 clusters, depending on the proteins’ expression in the wildtype (Fig 3C): ventral (eg. *snail*), lateral (*sog*), dorsal (*dpp*), dorsal and lateral (*grh or std*), dorsal and ventral (*ama*) or lateral and ventral (*neur*). However, in addition to these clusters (marked as 1/D, 2/L, 5/V, 6/DL,/DV and 12/LV in Fig. 3B,C, Supplementary Table 6) the clustering yielded a further 8 clusters (Fig. 3B,C). This results from the surprising difference between the two ventralising genotypes (Fig. 3B,C,E).

Most of the marker proteins above were found in their proper predicted class (Fig. 3D). Among the few allocated to clusters where the two ventral mutants differed in their behaviour, there was no general rule as to which of the two ventral mutants represented the ‘correct’ value. For example, for Heartless and Net, both of which are expressed in the mesoderm and also on the dorsal side of the embryo, one was seen with increased abundance only in *serpin27A* embryos, the other only in *Toll^10B^* embryos (Fig. 3D). Similarly, for genes that are excluded from the mesoderm, i.e. expressed in dorsal and lateral regions, some scored as present in lower abundance in *serpin27A* (e.g. crb), whereas others were reduced in *Toll^10B^*(eg. numb). We will return to the difference between *Toll^10B^* and *spn27A* below.

### 5) Comparison of RNA and protein expression patterns

Protein levels can be regulated post-translationally, and RNA and protein expression levels do not necessarily correlate strongly during development [23]. However, the regional distribution of proteins in the early *Drosophila* embryo is thought to be achieved mainly through transcriptional regulation that sets up the patterning of the AP and DV axes of the embryo [24, 25]. We therefore investigated how well the proteomes reflected known dorso-ventral modulation of gene expression.

We first looked for genes whose RNA expression patterns are reported in BDGP [26–28] *in situ*, (https://insitu.fruitfly.org/cgi-bin/ex/insitu.pl) to make a comparison between those with ventral expression in this data set and ours. We extracted all those genes that carry the labels ‘mesoderm’, ‘trunk mesoderm’, or ‘head mesoderm’ in BDGP (which are not mutually exclusive). 107 of the resulting set of 109 genes had their proteins detected in our analyses, and 71 had been allocated to one of the DV clusters. 60 were found in clusters that were fully or partially consistent with the reported RNA pattern (Supplementary Table 7). Of the 11 proteins among these 71 that show consistent mesodermal upregulation in both ventralizing mutants (DV cluster 5), all are reported as ventrally expressed in BDGP.

There is also a database representing an atlas of differential gene expression at single cell resolution for precisely the time window of early gastrulation [29] against which we compared the proteomes to regional RNA expression. Filtering out ubiquitously expressed genes left 8924 differentially expressed genes of which 3086 coded for 3120 proteins in our clustered proteome dataset (Fig. 4B).

**Figure 4.**
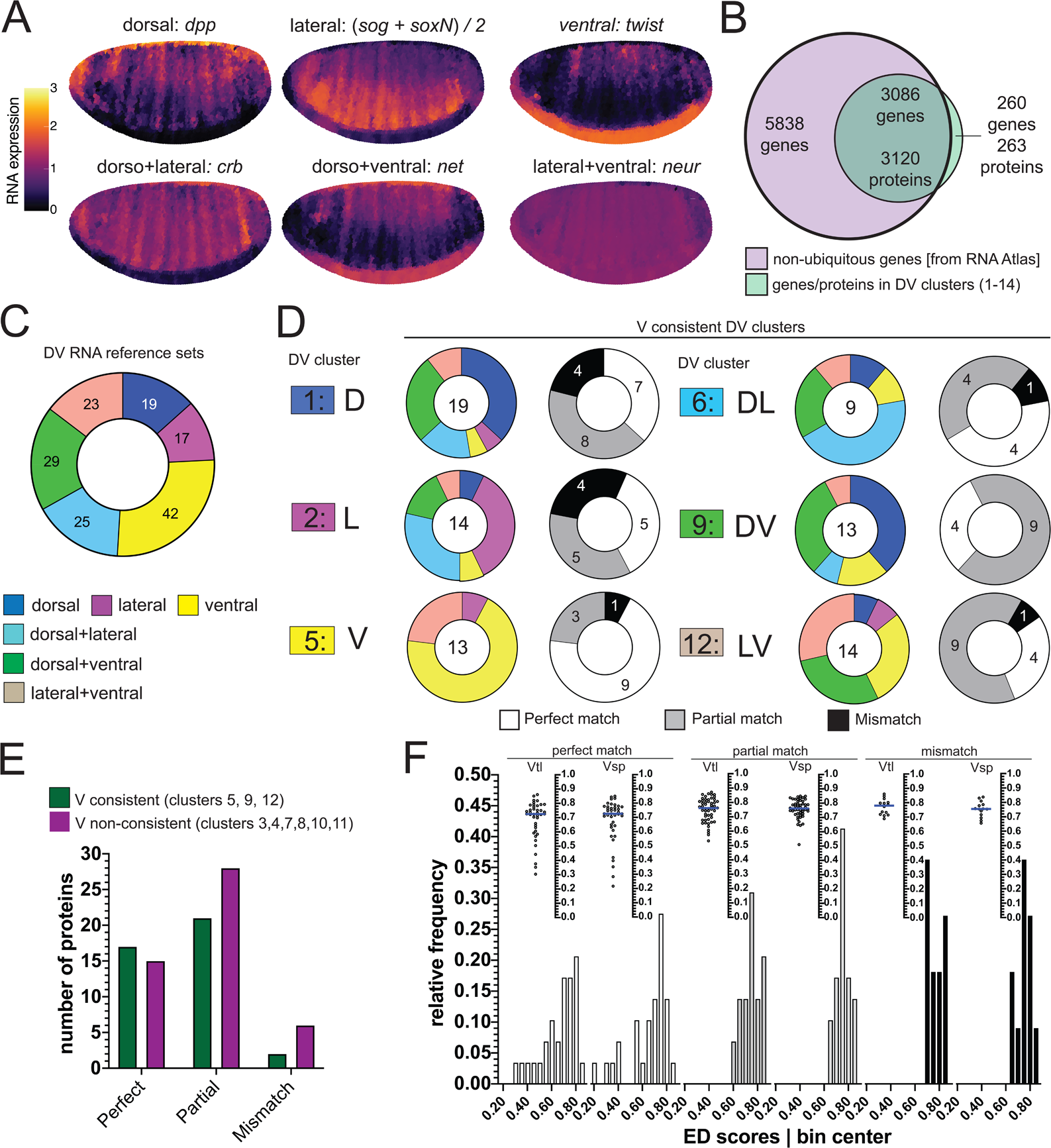
Comparison of RNA and protein expression patterns. **A)** Expression patterns of the reference genes used for assembling the RNA reference sets. Scale bar indicates RNA expression strength. Yellow depicts high RNA expression. **B)** Venn diagram showing the intersection between the genes of the RNA atlas with non-ubiquitous expression (8924) and the genes that encode the proteins assigned to DV clusters (3383/3393 proteins were successfully matched to a FBgn gene identifier). **C)** Number of non-ubiquitous genes allocated to each DV RNA reference set that trespassed the corresponding filter/threshold values (155/8924 genes, see methods and Supp. Fig. 4B-D). **D)** Matching of genes in the RNA reference sets with the proteome DV clusters in which the ventralized mutants display a consistent behaviour against the wild type (DV clusters 1, 2, 5, 6, 9 and 12, see Fig. 3B,C). Coloured pie charts represent the allocation of genes within each DV cluster to the RNA reference sets (colour code as in fig. 3). Grayscale pie charts represent the outcome of the comparison for filtered genes: white: perfect for an exact match, grey: partial, if the RNA reference included the correct match but also another region, black: mismatch, where the protein expression did not overlap with the RNA reference. Values in the centre of pie charts indicate the number of genes compared. **E)** Number of proteins in each outcome of the RNA-proteome comparison (perfect, partial, mismatch) in ventralized-consistent clusters (dark green) and ventralized non-consistent clusters (dark magenta) that belong to the same regulation categories: ventral (3,4,5), dorsal+ventral (7,8,9) and lateral+ventral (10,11,12). **F)** Two different representations, histogram and a swarm plot of the distributions of the Euclidean distance score for proteins that had a perfect, partial or mismatching overlay with the DV RNA reference sets. The histogram and the scatter plots are shown separately for calculations using each ventralized genotype: Vtl = *Toll^10B^/def*; Vsp = *spn27A^ex^/def*. In the swarm plots, each dot is a protein.

We first sorted these genes according to their expression patterns into the categories used above (D, V, L, DV, DL, LV) by virtue of similarity in their expression to six reference genes (Fig. 4A, Supplementary Fig. 4A-C). In a second step, we excluded those that showed only spurious differences in expression along the DV axis, ending up with 155 genes with clear DV differences forming six DV RNA reference sets (Supplementary Table 8, Supplementary Fig. D, Fig. 4C).

We then compared our 14 protein clusters against these six RNA reference sets. We asked for each protein which RNA reference set contained its corresponding gene. Theoretically, if both classifications, i.e. the RNA reference set and the proteomes, were perfectly correct, then genes from a protein cluster should be included only in the corresponding RNA reference set. We found that the majority of RNAs had proteins in partially or fully matching clusters of the proteomes (Fig. 4D, Supplementary Fig. 4E). For example, 9 of the 13 proteins (69%) in cluster 5 (ventral-consistent) found their gene in the ‘twist’ similarity reference group (ventral; a perfect match: white in pie charts: Fig. 4D, Supplementary Fig. 4E). The next best matches (e.g. ventral plus lateral, instead of only ventral; a partial match, grey) were often also highly represented: 3 of the 4 remaining cluster 5 proteins found their gene in the ‘neur’ similarity reference group (lateral+ventral).

Thus, the majority of proteins had perfect or partial matches with the RNA expression, showing that two independent measurements of regional expression patterns arrive at the same allocation. This confirms in an unbiased manner that the hierarchical clustering successfully sorted the proteomes in the correct manner, further supporting the initial assumption that the mutant populations were representative of specific regions in the embryo.

### 6) Different effects of the *Toll^10B^* and *spn27A* mutations on dorsal gene expression

The difference between the results for the *Toll^10B^* and *spn27A* embryos was an unexpected and potentially biologically interesting discovery. We investigated whether the matching of the protein distributions to their RNA expression patterns could give us further biological insights.

We find that for those clusters in which *Toll^10B^* and *spn27A* agree, a larger proportion of proteins is allocated to the correct RNA reference set than in the clusters in which *Toll^10B^* and *spn27A* differ (Fig. 4D,E, Supplementary Fig. 4E). The V cluster 5, in which *Toll^10B^* and *spn27A* agreed, included not only Snail and Twist but also other genes expressed in the mesoderm (Supplementary Fig. 2B), such as Mdr49 [11], CG4500 [8] and Traf4 [30].

For the proteins from the ‘ventral inconsistent’ clusters we found that the *Toll^10B^* mutant differs from the *spn27A* mutant in a consistent manner. Proteins classified on the basis of being upregulated in *Toll^10B^* (clusters 3, 7, 10 and 13) are often mismatched to genes with an ectodermal expression (dorsal and/or lateral RNA), whereas this does not occur for those classified based on their upregulation in *spn27A* (clusters 4, 11 and 14, Supplementary Fig. 4E). This means that although the morphological ventralisation and upregulation of ventral genes is strong in *Toll^10B^* mutants, the ectopic Toll signalling in the mutant fails to suppress all dorsal markers. This confirms previous suggestions that *spn27A* mutants retain no or almost no DV polarity whereas *Toll^10B^* embryos retain residual polarity [6, 31] and, as we now show, expression of many dorsal-specific proteins. Determining the developmental source of these differences goes beyond the scope of this study, but will warrant further investigation.

### 7) RNA-protein match versus degree of differential expression

We wondered whether there were consistent differences between those proteins that matched their RNA and those that did not. For example, a protein with large fold-changes may be more likely to match the correct RNA distribution. Because the clustering assigned proteins only on direction and not on extent of change, clusters also contain proteins with very small differences between the DV populations, even in cases where the RNA is known to show a clear difference (eg. Traf4; Supplementary Fig. 2C).

To distinguish between strong and weak differential expression, we ranked proteins by comparing them to the protein in each cluster that showed the greatest fold changes in the mutants over wildtype. We calculated the Euclidian distance (ED) between each protein and the protein with the greatest changes in a three-dimensional space where each axis represented fold-changes in one genotype (see methods, Supplementary Table 10). Thus, proteins with the lowest ED scores are those that are closest to the most extreme protein. We then analysed if this score correlated with the degree to which a protein matched the RNA expression of the corresponding gene. We found that proteins from the ‘matching’ groups had ED-scores that were skewed towards smaller values than those with non-matching proteins (Fig. 4F) indicating that proteins with more extreme expression differences (low ED scores) are more likely to match the correct RNA expression pattern.

In summary, these approaches stratify our results in a useful manner: first, the DV clusters in which the two ventralized mutants behave consistently represent better the RNA expression patterns; second, proteins with strong fold-changes are more likely to represent the distribution of the corresponding RNA.

### 8) The phosphoproteome of embryonic cell populations during gastrulation

Changes in the abundance of phosphosites may occur for two reasons: either the protein itself varies in abundance, or the protein level is constant, but the protein is differentially phosphorylated. Combinations of these cases are possible, and protein abundance may be affected by phosphorylation itself. Since we know the changes in protein abundance, we can distinguish these cases by comparing the full proteome against the phospho-proteome (with the caveat that, for technical reasons, our measurements were done on parallel experiments rather than on the identical samples).

1765 of the phospho-proteins (96%) we identified were ones that we also found in the proteome, whereas 82 had not been detected in the proteome (Fig. 5A). The match between the proteome and phosphoproteome yielded 6297 pairs of phosphosites and their host proteins across all genotypes. We tested for each differentially phosphorylated site whether its respective protein was up- or down-regulated and made this comparison for each mutant genotype versus the wildtype. For the proteome we applied the same threshold as used above for clustering (+/-1.4 fold change), and for the phosphoproteome we used +/-1.3 fold change (see methods). We found that most of the changes in phosphorylation were in proteins for which the level of the protein itself was unchanged (black and white boxes in Fig. 5B; 67 to 82% of the protein-phosphosite pairs). Among those for which the host protein showed differential abundance, 7 - 13% of their phosphosites changed in the same direction (both protein abundance and phosphorylation up, or both down), and 10 - 19% of their phosphosites changed in the opposite direction.

**Figure 5.**
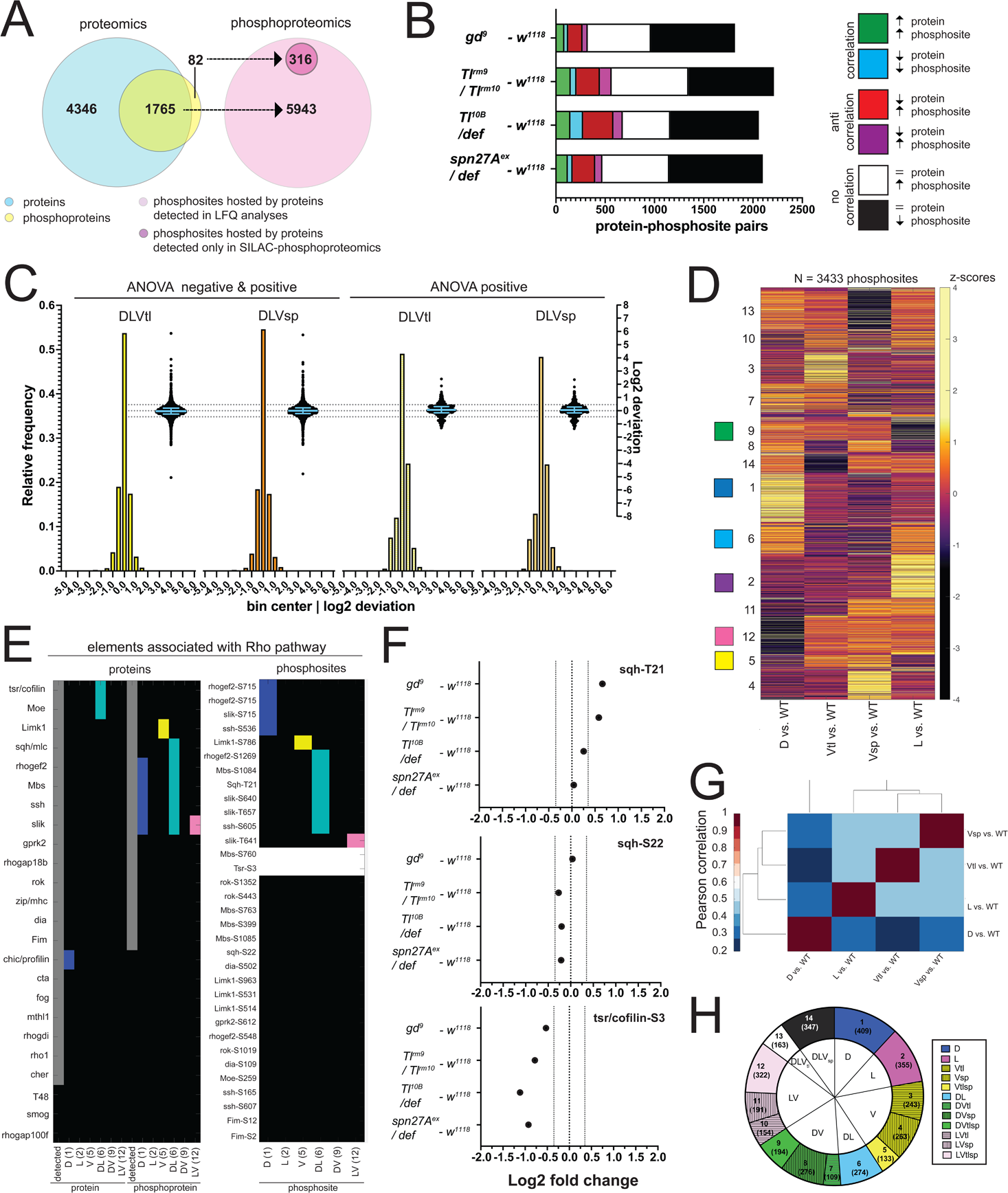
The phosphoproteomes of the mutant embryos. **A)** Match between detected protein groups in the proteomic and phosphoproteomic experiments. Left: blue: protein groups detected in proteomes (LFQ); light green: protein groups detected both in proteomic and phosphoproteomic experiments; yellow: protein groups not detected in proteomes but detected in phosphoproteomes. Right: pink: phosphosites hosted by a protein group detected in the proteomic analyses; magenta, phosphosites that could not be matched to a protein group detected in the proteomes. **B)** Correlation between the fold changes (FCs) of phosphosites and their host proteins in DV mutants vs. wild type. Correlations: FC of protein and phosphosite are both positive (green) or negative (blue). Anti-correlation: FC of protein and phosphosite have different signs (red and magenta). No correlation: protein levels are unchanged but phosphosite FC is positive (white) or negative (black). Bars represent the number of phosphosite-host protein pairs falling in each correlation category within each DV mutant vs. wild type comparison. **C)** Two different representations, a histogram and a swarm plot, of the deviation parameter (in log2 scale) calculated for each of the two ventralized genotypes (D: dorsalized, L: lateralized, Vtl: *Toll^10B^*/*def*, Vsp: *spn27A^ex^*/*def*). This was done once for all phosphosites present in all genotypes, and once only for those that were ANOVA positive. In the swarm plot, each dot represents a phosphosite. The y-axis for the histograms is shown on the left, for swarm plots on the right. Blue bars show the median with inter-quartile range (IQR). The median is close to zero and the IQRs range from −0.2387 to 0.3197. The dotted line indicates the 0.35 and −0.35 deviation in swarn plots. Histograms and swarm plots were assembled with phosphosites detected in all genotypes. **D)** Clustergram of the hierarchical clustering of 3433 phosphosites. Z-scores were calculated using the thresholded fold changes between mutants and wild type. Coloured boxes indicate the clusters with consistent behaviour for the two ventralising genotypes (for colour-coding see Fig. 3D). Numbers identify the different clusters for reference across panels, and are equivalent to the proteome (Fig. 3B,C). **E)** Detection and predicted regulation (DV clusters) of Rho pathway proteins and phosphoproteins (left panel) and the corresponding phosphosites (right panel). Colours mark proteins, phosphoproteins or phosphosites in DV clusters with ventralized consistent behaviour(DV clusters 1, 2, 5, 6, 9 and 12). Grey boxes represent the detection of a particular protein or phosphoprotein in the wild type genotype. White boxes represent an increased or decreased abundance in all DV mutants vs. wild type. **F)** Log2 fold changes (FC) of known phosphosites in sqh: T21 (top) and S22 (center) and tsr/cofilin S3 (bottom). Dotted lines indicate log2 FC = 0, 0.35 and −0.35. Colors depict DV mutant genotypes and their corresponding comparisons against wild type: blue: dorsalized, magenta: lateralized, yellow: ventralized (*Tl^10B^*/def and *spn27A^ex^*/def). Bars depict mean and standard error of the mean across replicates. Absence of a dot indicates the protein was not detected in a particular condition or log2 FC calculation not feasible, absence of error bars in Log2 intensity indicate protein was detected in a single replicate. Dotted line indicates log2 FC = 0, log2 FC = 0.5 (for proteins) or log2 FC = 0.35 (for phosphosites). **G)** Correlation matrices using Pearson correlation coefficient between the fold changes of each mutant vs. wild type comparison in the phosphoproteome experiment. **H)** Pie chart showing the number of phosphosites allocated to each DV class in (C).

We first tested if the phosphoproteomes fitted the ‘linear model’ (i.e. whether the sum of the weighted mutant values corresponded to the measured values in the wildtype). We found that the majority of the phosphosites indeed fitted this model (Fig. 5C, Supplementary Table 5). Among the strongly deviating phosphoproteins, we find a number of kinases with known morphogenetic functions, such as Par-1, SRC42A and nucleoside-diphosphate kinase (awd) have known morphogenetic functions (Supplementary Table 12 and Supplementary Fig. 3E).

We clustered the phosphosites using the same procedure as for the proteome (see methods). Thresholding excluded 1234/6259 sites defined as unchanged across all comparisons against wild type. We also excluded those sites that were either upregulated or downregulated versus the wildtype in all mutants (615/6259) and clustered the remaining 3433 phosphosites, again yielding 14 DV clusters (Fig. 5D,H, Supplementary Table 6). The two ventralising mutants now clustered together, and the dorsalized mutant showed the most distinct behaviour (Fig. 5G).

### 9) Enrichment of cytoskeletal components among differentially phosphorylated proteins

One aim of this study was to find cellular components that are differentially modified along the DV axis and that are candidates for regulating cell shape. Most likely, these cellular components are regulated by protein complexes or interacting protein networks, as already known for the regulation of actomyosin by the Rho pathway and some components of adherens junctions.

Signalling through the Rho pathway provides an important mechanism involved in morphogenesis. Rho is activated and necessary for cell shape changes in the mesoderm, but we do not know the full set of the components of the pathway that are modulated in the mesoderm or elsewhere along the DV axis. Of 24 proteins associated with Rho signalling, we detected 21 in the wild type and at least one of the mutants (Fig. 5E). Most occurred at similar levels in all genotypes (including the myosin light chain), except Twinstar/Cofilin, Moesin and chickadee/Profilin, which were present in increased abundance in the ectodermal cell populations (clusters D and DL).

14 of the 21 proteins were phosphorylated. The detected sites included the known phosphosites at Threonine-21 and Serine-22 in the myosin light chain, and at Serine-3 in Cofilin/Twinstar (Fig. 5F). Six of the 14 phosphorylated proteins had at least one site that was differentially phosphorylated along the DV axis (Fig. 5F). In addition to Cofilin/Twinstar, the phosphorylation state of both its regulatory kinase LIMK1 and its phosphatase Slingshot (ssh) were modulated in the D and DL clusters. Myosin light chain showed increased phosphorylation at T21 in the DL cluster (Fig. 5E,F), and RhoGEF2 and the myosin phosphatase Mbs were differentially phosphorylated in the DL and D clusters (Fig. 5E). In summary, we detected most of the elements of a well established pathway required for gastrulation and also identified new candidate regulation nodes within the Rho pathway.

To find such networks, we used a diffusion-based algorithm [32] on each of the DV clusters. The starting weight of each protein was [based on its ED score and deviation from the linear model. Since these scores existed separately for the two ventralizing mutants, we also had to conduct the analyses twice in each case, i.e once for each dataset. We focussed our further analyses only on those proteins that behaved in the same way for the two ventralising genotypes.

Overall, we generated 24 protein networks for the proteome and 24 for the phosphoproteome, i.e. for each of the six DV clusters (D(1), L(2), V(5), DL(6), DV(9) and LV(12)) carried out with four different scores. The significance for each of the 48 networks was tested by comparing the values of the nodes against those obtained in 1000 random networks. Only those nodes with p-values <0.05 were included in the final networks. These networks reflect the global effect of the change in each protein on its network neighbours. To further distil the networks to the ‘active’ signatures we then investigated the local impact of this effect using an ego network analysis (see methods). The ego network analysis extracts for each seed protein those neighbours that are most similar in function and most interconnected to the seed. These nodes represent the foreground for the enrichment analysis of the cellular components, yielding a set of 83 and 87 ontology terms that were enriched for at least one condition in the proteome or phosphoproteome, respectively.

We concentrated our further analyses only on those ontology terms that were enriched for at least two of the four scores, (ED score or deviation, each for either of the ventralized genotypes for any of the DV clusters; proteome: 63, phosphoproteome: 62 cellular component terms), and used a heatmap to represent them (Fig. 6A,B and Supplementary Fig. 5A,B). The heatmaps illustrate that most of the enriched cellular components in both experiments were associated with DNA and RNA metabolism or the regulation of gene expression, which is not unexpected for this developmental period of dynamic changes in gene expression. In agreement with this, the majority of the enriched proteins and phosphoproteins were characterised as nuclear ontology classes. Because of our interest in morphogenesis we focused on the cellular components that belong to cytoskeletal, cell adhesion and vesicle trafficking categories. In the phosphoproteomes the networks enriched for cytoskeletal components were much more prevalent in the phosphoproteomes (14 of 63) than in the proteomes (3 of 62), with microtubules strongly represented (12 of 14 cellular components), in particular the alpha and beta tubulins and microtubule associated proteins such as the Kinesin Heavy Chain, and centrosomal proteins (Supplementary Table 11). Cytoskeletal proteins are often localised in the cell cortex, and we indeed find this association reflected in the results of the network analysis. The cell cortex is among the enriched components, and among the proteins in this category, we find cytoskeletal elements. For example, networks that include the actin-microtubule crosslinker Shot and the actin polymerase Profilin are enriched in the dorsal cluster; networks that include the apical polarity determinant Stardust or the Hippo pathway component Warts in the dorso-lateral cluster. A phosphoprotein network associated with adherens junctions and zonula adherens one of which contains the junction-actin connectors Canoe and Girdin [33, 34]was enriched in the D cluster (Supplementary Table 11).

**Figure 6.**
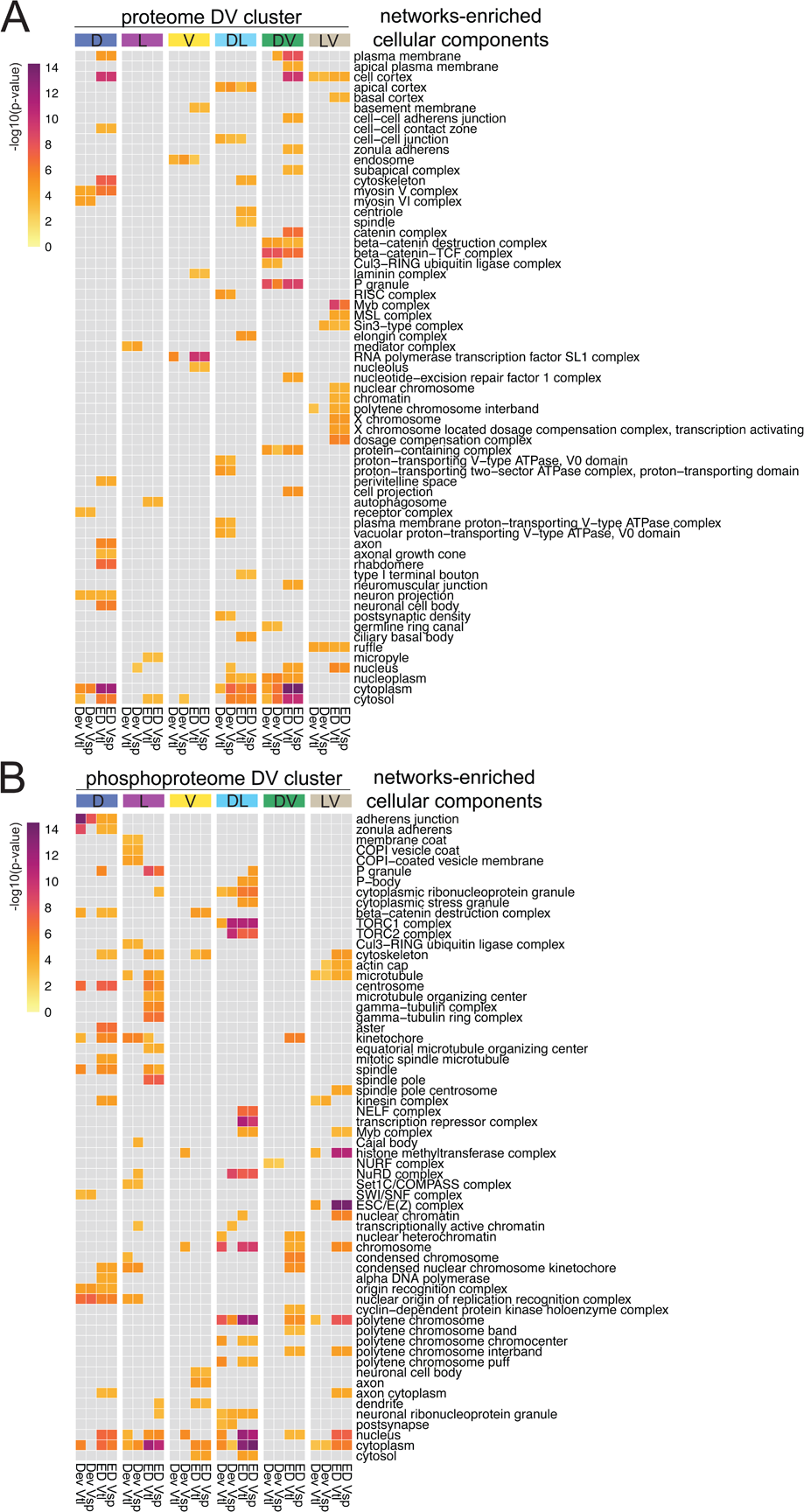
Diffused network analyses of DV proteomes and phosphoproteomes. Heatmap representations of significantly enriched cellular component ontology terms in at least two networks across all DV clusters with a consistent behaviour in their ventralized genotypes (DV clusters 1, 2, 5, 6, 9 and 12). **A)** Cellular components enriched in networks emerging from the proteome. **B)** Cellular components enriched in networks emerging from the phosphoproteome. Scalebar indicates the -log10(p-value) for a measure of statistical significance across ontology terms and DV clusters. For each DV class, four diffused networks were generated using the deviation (’Dev’), and the euclidean distances (’ED’) to score the nodes of emerging networks. Calculations were performed independently for each score and each ventralized genotype: Vtl (*Toll^10B^/def*) and Vsp (*spn27A^ex^/def*).

In summary, we can highlight two outcomes of the network propagation analysis. First, most networks, whether derived from the proteomes or the phosphoproteomes, are enriched for cellular components associated with regulation of gene expression (transcription, epigenetic regulation, translation, protein turnover). This is a useful validation of the approach, in that it reflects the main biological process that occurs at this stage of development: giving cells in the body different developmental fates, which is achieved through setting up different gene expression programmes.

Secondly, the cytoskeleton emerges as a major target of regulation, especially in the phosphoproteome, with the most prominent component being the microtubules. This is an interesting target for further exploration in the context of gastrulation and fits well with recent results that microtubules play a role in epithelial morphogenesis [35–38].

### 10) Functional implications of networks enriched for microtubule components

To test the biological relevance of these findings, we chose microtubules (MTs) as an example to study functionally in the embryo. Before gastrulation, all cells along the DV axis have two subpopulations of MTs: a disordered apical network of non-centrosomal MTs with short, non-aligned filaments, and an ‘inverted basket’ of basal-lateral MTs originating from the centrosomes and enclosing the nucleus [35, 39] (Fig. 7A). MT dynamics correlate with acetylation (stable MTs) and tyrosination (dynamic MTs) of α-tubulin [40]. We explored in fixed embryos markers for MT dynamics along the DV axis. We detect acetylated α-tubulin only among the basal-lateral MTs (Fig. 7A, Supplementary Fig. 6A, blue arrows) whereas tyrosinated alpha-tubulin is seen in both MT populations (Supplementary Fig. 6A, arrowheads; see also Takeda et al 2018 [35]).

**Figure 7.**
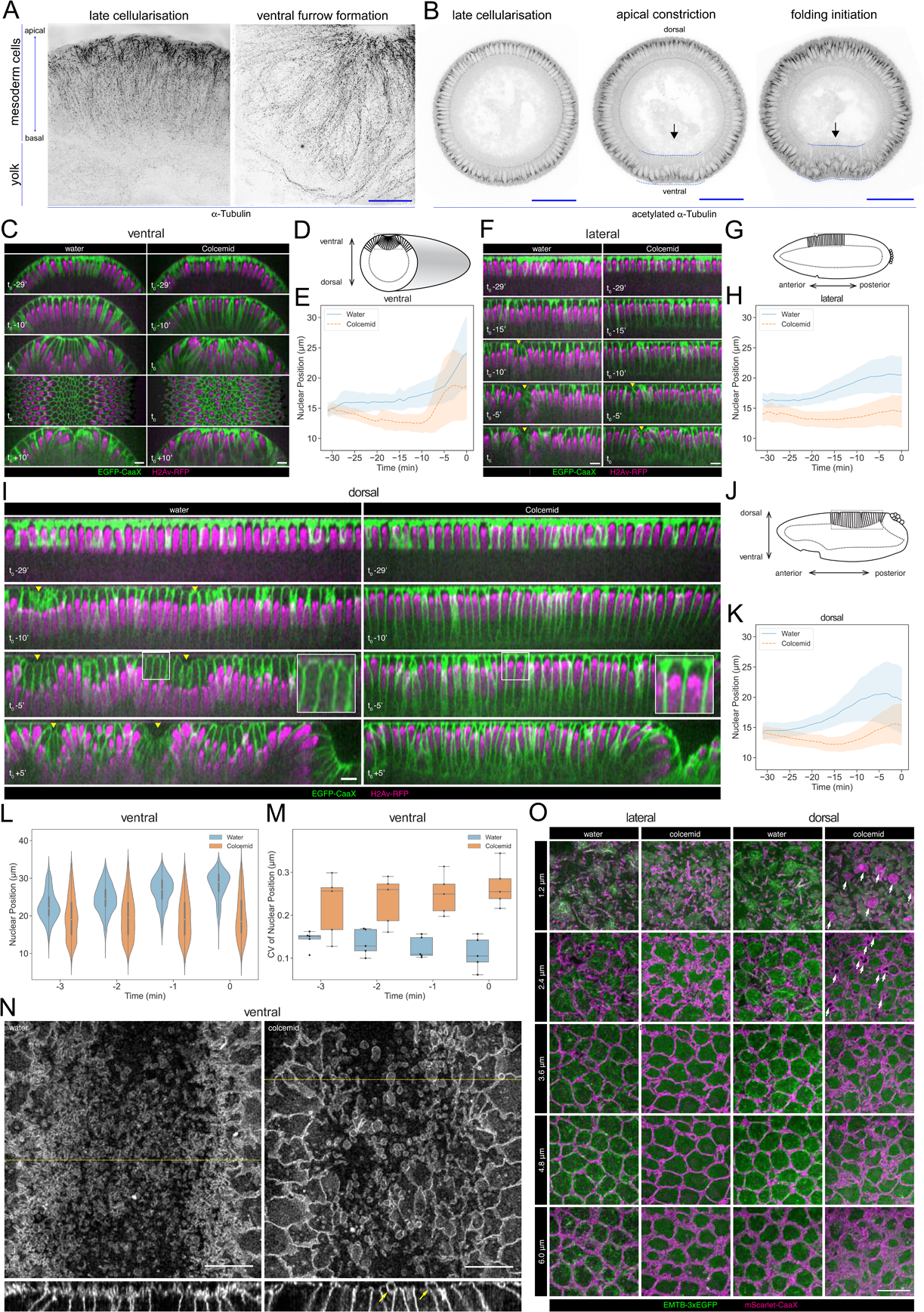
Microtubule organisation and *in vivo* functions. **A)** Images (OMX super-resolution microscope, max-projected) of mesoderm cells (ventral domain) using physical cross-sections from fixed embryos stained with antibodies against α- Tubulin. Left panel: onset of gastrulation, right panel: contractile mesoderm during ventral furrow formation. Scale bar is 10 μm. **B)** Images (confocal, max-projected) of physical cross-sections from fixed embryos stained with an antibody against acetylated α-Tubulin at the onset (left panel) and during ventral furrow formation (center panel: initiation of gastrulation, apical constriction; right panel: mesoderm folding). Arrow indicates detection of acetylated α-Tubulin specifically in basal-lateral microtubules (inverted basket). Dotted blue line encloses mesodermal cells, in which a progressive reduction of acetylated α-tubulin is detected during ventral furrow formation. Scale bar is 50 μm. **(C, D, E)** Phenotypic effect of colcemid injection on the ventral side of the embryo during cellularization and early gastrulation. **C)** Time-lapse series of Z re-slice (and a surface projection for the t_0_ time point) showing cellular and tissue architecture with membrane (EGFP-CaaX) and nucleus (H2Av-RFP) labels. **D)** A schematic drawing of tissue architecture during ventral furrow formation with a dotted rectangular box depicting the ROI of the re-slice view in panel C. **E)** Nuclear position, i.e. distance from the embryo surface, as a function of time during cellularization. **(F, G, H)** Same as above for the lateral side of the embryo with **F)** time-lapse series of Z re-slice, **G)** a schematic drawing during cephalic furrow formation and a dotted rectangular box for the ROI in panel F, and **H)** nuclear position as a function of time. **(I, J, K)** Same as above for the dorsal side of the embryo with **I)** time-lapse series of Z re-slice, **J)** a schematic drawing during dorsal fold formation and a dotted rectangular box for the ROI in panel E, and **(K)** nuclear position as a function of time. Insets, enlarged view showing the shape of the apical dome. **(L, M)** Colcemid treatment leads to a wider distribution of nuclear positions in apically constricting VF cells during the early phase of apical constriction, shown in a violin plot for nuclear centroid position **(L)** and a box plot for its coefficient of variation **(M)**. For all of the above, t_0_ represents the onset of gastrulation, as defined in M&M. Yellow arrowheads: surface clefts resulting from cephalic furrow **(F)** and dorsal fold **(I)** initiation. **(N)** Apical surface projection (top row) of membrane (3xmScarlet-CaaX) and Z re-slice (bottom row; taken from the yellow dotted lines in the top row) showing enlarged membrane blebs (yellow arrows) after colcemid injection during ventral furrow apical constriction. **(O)** Apical membrane phenotypes in the lateral and dorsal cells observed at different Z positions, each with a 1.2 μm projection, visualised with membrane (3xmScarlet-CaaX) and microtubule (EMTB-3xEGFP) labels. White arrows: abnormal membrane blebs that are devoid of microtubules and observed exclusively on the dorsal side. Scale bars: 10 μm.

When the ventral furrow begins to form, we detect differential acetylation of the basal-lateral MTs along the DV axis. In the ectoderm, MTs become increasingly acetylated but retain their original organization (Fig. 7B, Supplementary Fig. 6A). In constricting mesodermal cells, the basal-lateral MTs become less acetylated as they move basally together with the nuclei (Fig. 7B, cells along dotted line, Supplementary Fig. 6A). MTs in non-constricting mesodermal cells next to the furrow bend in the direction of cell stretching as the apical surfaces of these cells extend towards the constricting mesodermal cells at the ventral midline (Supplementary Fig. 6B, arrow). These MTs are non-acetylated, but partially tyrosinated (Supplementary Fig. 6A, blue arrowhead), and are therefore probably newly polymerized.

We depolymerised microtubules with colcemid and observed the consequences for cellular dynamics prior to and during epithelial fold formation on the ventral, lateral and dorsal sides of the embryo. To prevent cellularisation defects [41], we injected colcemid when the cellularising front had reached a depth of 15∼20 μm. Less than 1 min after the injection, most apical filamentous structures, astral MTs emanating from the centrosome, and the centrosomes themselves disappeared. MTs associated with the nuclear envelope were retained (Supplementary Fig. 6C,D), consistent with the presence of acetylated MTs in the inverted basket structure, which likely resist depolymerization by colcemid.

In control embryos, nuclei moved basally for 1∼2 μm in the last ∼20 minutes of cellularisation on all three sides of the embryo. In embryos injected with colcemid, nuclei moved slightly in the opposite direction towards the very apical end of the cell (Fig. 7C-K, Supplementary Movies 1-3).

During ventral furrow formation, nuclei moved basally in the constricting cells, creating a large gap between the nucleus and the apical surface of the cell. In colcemid-injected embryos, nuclear positioning was more randomised (Fig. 7C, t_0_; 7L,7M). Ultimately, the ventral furrow failed to form (Fig. 7C, t_0_+10’, Supplementary Movie 1).

On the lateral side of the embryo, a single or a double row of cells initiate the formation of the cephalic furrow through lateral actomyosin contraction [42]. In colcemid-injected embryos, the cephalic furrow forms normally, but with a delay of ∼5 min although nuclei are also apically positioned on the lateral side (Fig. 7F-H, Supplementary Movie 2).

On the dorsal side, the formation of dorsal folds depends on the remodelling of apical MTs, but not on myosin contractility [2, 35, 43]. All dorsal cells form an apical dome with a characteristic, curved cell apex prior to the initiation of dorsal folds (Fig. 7I, t_0_-5’, insets), contrasting with the flat apical surfaces of ventral and lateral cells during late cellularization. In colcemid-injected embryos, the apical surfaces of dorsal cells remain flat, supporting the model that MT-dependent force is required for apical dome formation (Fig. 7I, t_0_-5’, insets). At the onset of dorsal fold formation, the apical dome of the initiating cells descends from the embryo surface as a result of cell shortening, producing a cleft (Fig. 7I, t_0_+5’). Dome descent does not occur in the colcemid-injected embryos, and dorsal fold formation eventually fails (Fig. 7I, t_0_+5’, Supplementary Movie 3), supporting the current model that differential microtubule forces underly localised cell shortening, that initiates dorsal fold formation [35, 43]. In sum, depolymerization of MTs reveals distinct functional requirements of MTs in all three types of epithelial folds in the gastrulating embryo.

We next examined the effect of MT depolymerization on the apical plasma membrane. During ventral furrow formation, the apical membrane of the constricting cells produces numerous membrane blebs that result from a transient detachment of apical membrane from the contracting actin cortex [44]. Colcemid injection yields enlarged membrane blebs in constricting ventral furrow cells (Fig. 7N). Lateral and dorsal cells lack these constriction-dependent blebs. Instead, they accumulate excessive apical membrane, displaying tortuous subapical membrane just prior to the onset of gastrulation (Fig. 7O) [45, 46]. After colcemid injection, we also observed a distinct class of micron-scale membrane blebs in all dorsal cells, not limited to the dorsal fold initiating cells and unrelated to myosin-dependent apical constriction (Fig. 7O). These blebs form during mid to late cellularization, exclude MTs and are stable for minutes. The mechanistic basis of this new class of blebs would require further investigation.

In sum, as predicted by diffused network analyses, all cell populations use microtubules during gastrulation. Microtubules are required for correct nuclear positioning and cell shape homeostasis. Distinct phenotypic consequences of the apical membrane following colcemid injection suggest differential functionality in the maintenance of membrane-cortex attachment or the dynamics of apical membrane retrieval for MT networks residing on different sides of the embryo.

## Discussion

We have presented a large-scale study of regional differences in the proteome of the early *Drosophila* embryo. We looked at a stage soon after the maternal to zygotic transition in gene expression, namely the onset of morphogenesis. We can compare our results to a previous study on regional differences in the proteome at this stage that used mutants, as we did, to represent different regions of the embryo, and in that regard should be directly comparable. This study was based on 2D gel electrophoresis combined with mass spectrometry, which, while ground-breaking at the time, allowed only a small number (37) of unique proteins to be identified. All of these were also detected in our proteomes.

Because the differential detection in this study was based on PAGE it was possible to defect different protein isoforms and therefore differences that may be due to phosphorylation. Of the proteins with variable isoforms, we found that 15 were phosphorylated in our own study, of which seven show differences in the mutants, and all of these are consistent with the changes seen in the 2D-PAGE experiment [10].

Comparing protein abundance against RNA expression could, in principle, reveal which proteins are post-transcriptionally regulated, but this can only be done if the techniques and approaches are as near-identical as possible, and if the results are technically perfect. Thus, even comparing differential RNA expression data obtained with different methods yields only partially overlapping results. For example, an Affymetrix-based study that again used mutants to represent regions along the DV axis of the embryo [11] identified 23 genes for which the RNA levels were higher in ventralized than in lateralized or dorsalized embryos. Comparing those to the expression patterns determined by single-cell RNA sequencing [29] reveals that five appear to have little or no dorso-ventral modulation, a result that is also confirmed in the BDGP in situ hybridisation database. Those genes previously identified by genetic or functional studies, and known to be involved in mesoderm development (including marker genes like twist, snail, zfh1, htl etc) show up in all studies.

Thus, a comparison of our proteome data to reported RNA expression patterns has to be seen with caution. Nevertheless, such comparisons showed good matches for the abundant, well-studied genes and proteins: We detect the proteins for 13 of the 17 genes that are seen as ventrally upregulated genes in both studies [11, 29]. Of those, we see all but four as ventrally upregulated, again including known ventral marker genes.

These comparisons lead to the question of how to judge which of the differences in protein abundance or regulation are biologically relevant and therefore interesting to follow up with functional studies. Confining the selection to those that are consistent with other studies would defeat the purpose of the experiment. Similarly, choosing the extent of change as a threshold would also exclude proteins we know to play a role in morphogenesis at this stage but which show only very small differences in expression. One example is Traf4 [30], which is active in the mesoderm, but expressed there at low levels, and becomes expressed in the ectoderm as gastrulation begins. In our experiment, it was strongly downregulated in the dorsalized embryo, but showed only sub-threshold upregulation in the ventralized embryos.

To obtain a better picture of processes or cellular components involved in the functional differentiation of the cell populations, rather than looking at individual genes, we identified networks of functionally related proteins that were enriched among the differentially regulated entities. We would like to highlight here the mechanisms of differential protein degradation, mRNA regulation and microtubule modifications.

A role for protein degradation in creating differential functions along the DV axis has previously been illustrated by the case of the E3-ubiquitin ligase Neuralized (Neur) which is required and upregulated in the prospective mesoderm [47]. The network analysis identified the cullin complex as differentially expressed and differentially regulated. We also find Neur in increased abundance ventrally. Known biological data thus validate the relevance of this network, which may in turn help to identify the as yet unknown targets for Neur in the mesoderm.

Another mechanism for post-transcriptional gene regulation is the differential translation or degradation of mRNAs along the dorso-ventral axis, and we find an enrichment of P-granule-related networks both in the proteome and the phosphoproteome. These networks are enriched within DV clusters with complete or partial ectodermal fate, i.e. the same clusters that show a strong uncoupling between mRNA and protein abundance. Partial agreement between mRNA and protein spatial distribution is not an exclusive feature of the gastrula: it has also been described for larval tissues derived from the ectoderm and neuroectoderm, where nearly all studied genes show mRNA/protein discordance) [48] (97.5%; N = 200 proteins). Therefore, the uncoupling between mRNA and protein abundance seems to be the rule rather than exception in at least these tissues, highlighting the importance of post-transcriptional regulation on gene expression regulation during development.

The diffused networks also showed phosphorylation of microtubules as a differentiating mechanism along the dorso-ventral axis during gastrulation, an interesting finding, because in epithelial tissues microtubules are often required for cell shape homeostasis [49] [50]. Morphogenetic cell shape changes in *Drosophila* for which microtubules are essential include the squamous morphogenesis of the amnioserosa [38] and the invagination of the mesoderm [36] and the salivary placode [37]. Here, we found that dorsal fold formation also requires microtubules. Ventral furrow and dorsal fold formation differ in their dependency on myosin [7], but our results show that both require microtubules for the basal relocalisation of nuclei. This requirement is functionally distinct from the association of microtubules with actomyosin during myosin-dependent tissue folding [36, 37, 51] and instead, may relate to the classic role of microtubules in vectorial trafficking and organelle localisation [52, 53]. One reason why nuclei need to be actively repositioned may be that in their apical location they constitute a physical barrier to the cell’s apical constriction.

### The differential proteomes and phosphoproteomes of the Toll^10B^ and spn27A ventralising mutants

Both *Toll^10B^* and *spn27A* ventralising mutations produced embryos that recapitulated known biological qualities of the mesoderm along the entire DV axis, such as the expression of ventral fate determinants, or the apical localisation of the adherens junctions. However, these similarities were not fully mirrored in their proteomes. Curiously, the *spn27A* proteome seemed to be more similar to the dorsalized than to the *Toll^10B^* proteome which would indicate that *Toll^10B^* embryos are ‘more’ ventral than *spn27A* embryos. However, most of the mesodermal marker genes (*snail, twist, mdr49, wntD, neur*) make an exception are more abundant in *spn27A* embryos. Similarly ectodermal fate markers are more strongly downregulated in *spn27A* than in *Toll^10B^* embryos. Specifically, *Toll^10B^* mutants fail to repress the expression of ectodermal genes such as *egr, zen* and *crb*.

How can ventralising mutations that act on the same pathway yield different proteomes? Spn27A is a serine protease inhibitor of the pathway that creates the active form of Spätzle (Spz), the ligand for Toll. Both mutations lead to constitutive activity of Toll, *Toll^10B^* through a mutation in the receptor itself [54, 55], *spn27A* through enabling a homogeneously high level of Spz along the DV axis [31] (rather than a peak on the ventral side). Because Spz is highly abundant it should not be a limiting factor for the activation of Toll [56] [57] (Supplementary Fig. 2A) and loss of Spn27A should enable the full activation of Toll along the embryonic DV axis. Our results indicate that constitutively active Toll does not lead to the same level of signalling as the binding of the ligand to the receptor, and that these different levels lead to unexpected differences in the downstream targets of the signalling pathway.

### The biological significance of deviations from the linear model

The ‘linear model’ we formulated is based on the assumption that each mutant embryo faithfully represents one defined area of cells along the DV axis of the embryo, and that the full set of cell types in the embryo can therefore be reconstituted as the sum of the mutant cell types - weighted according to the area they occupy in the embryo. This should also be recapitulated for any individual protein expressed in the embryo. We found this to be true not only for the trivial cases of those proteins that occur at equal level in all genotypes, but also for most of the differentially modulated proteins. However, some proteins and phosphosites did not fit the model but showed strong deviations. One explanation could be that in the normal embryo the embryonic regions communicate with each other, and this communication is necessary for the expression or modification of certain proteins. These interactions cannot occur when the fates occur in isolation from each other in the mutants, and therefore some proteins would not be regulated properly and would not fit the model. Thus, wherever an interaction between the cell populations in the embryo is necessary for generating the correct expression or phosphorylation level, the linear model we proposed no longer applies; this means that strong deviations may indicate non-autonomous regulation. We do know some genes whose expression along the DV axis is determined by input from neighbouring regions, such as Sog, Ind and single-minded. We indeed find that one of those proteins, Ind, is an outlier (deviation = 2.6) with higher than predicted expression in the dorsalized and ventralized mutants, consistent with repressive input from these regions in the wildtype.

Another case of proteins not following the linear model are those that are found either in decreased or increased abundance in all genotypes, a behaviour we observed for a small percentage of proteins and phosphosites, perhaps as part of a general stress response related to the mutant situation. This is illustrated by the most extreme example, TM9SF4, which encodes an immune-related transporter that is present in all mutants at nearly 100-fold higher levels than in the wildtype. However, we did not find that this was a general rule either in the proteomes or in the phosphoproteomes: stress-related categories such as those from the ‘chaperone’ or ‘immune response’ ontology classes represented only a small percentage of proteins and phosphoproteins with the highest deviations (Supplementary Fig. 3D,E).

## Methods

### *Drosophila* genetics and embryo collections

*w^1118^* (wildtype/control genotype in our studies, Bloomington stock 3605), *gd^9^*/FM6a [58] (provided by S. Roth), *Tl^rm9^* and *Tl^rm10^* [59] (provided by A. Stathopoulos), *Toll^10B^* [54] [55], *Df(3R)ro80b*/*TM3* (Bloomington stock 2198), *spn27A^ex^*/*CyO* [31], (provided by S. Roth, Bloomington stock 3764), *Df(2L)BSC7*/*CyO* [31] (provided by S. Roth). To visualise non-muscle myosin *in vivo*, a sqh-sqh::mCherry transgene (Bloomington stock 59024) was used to construct the following stocks *gd^9^*;*sqh-sqh::mCherry*/*CyO*, *sqh-sqh::mcherry*/*CyO*;*Tl^rm9^* and *sqh-sqh::mcherry*/*CyO*;*Df(3R)ro80b*/*TM3*.

Dorsalized embryos were derived from *gd^9^* homozygous female mothers, lateralized embryos were derived from trans-heterozygous *Tl^rm9^*/*Tl^rm10^* mothers, ventralized embryos we derived from *Toll^10B^*/*Df(3R)ro80b* and *spn27A^ex^*/Df(2L)BSC7 mothers (see Supplementary Table 1). Female mutant mothers were crossed with *w^1118^* males, and the F1 from each of these crosses were collected and processed for mass-spectrometry analyses.

To visualise Myosin Light Chain, we generated the following mothers: Dorsalized: *gd^9^*;*sqh-sqh::mCherry*/*+,* Lateralized: *sqh-sqh::mcherry*/*+*;*Tl^rm9^*/*Tl^rm10^*, Ventralized: *sqh-sqh::mcherry*/*+*;*Toll^10B^*/*Df(3R)ro80b*. Female mutant mothers were then crossed with *w^1118^* males, and from F1 of these crosses, embryos in stage 5a,b [60] were hand-selected under a dissecting microscope and mounted for live imaging (see below).

#### Embryo collections

Embryos collected half an hour after egg-laying were allowed to develop for 2hs 30’ at 25°C in a light and humidity-controlled incubator and then dechorionated in 50% bleach for 1’ 30”, washed with H_2_0 and visually inspected under a dissecting microscope (Zeiss binocular) for 15’-20’ at RT to ensure younger embryos from each synchronised collection had progressed into the gastrulation stage (Stages 6a,b [60]). Any embryos that had developed beyond the initial stage of gastrulation (as judged by morphological criteria, Stage 7 [60]) were discarded and the remaining embryos were placed in 0.5 ml Eppendorf tubes and flash-frozen in liquid nitrogen. An 0.5ml Eppendorf tube filled with embryos yields approximately 1 mg of protein.

#### Transgenic fly lines generated in this work

Three transgenic lines were generated to visualise cell membranes and microtubules. For cell membranes, a single copy of EGFP or three copies of mScarlet interleaved with linkers (DELYKGGGGSGG) were trailed by a C-terminal CaaX sequence from human KRas4B (KKKKKKSKTKCVIM) for membrane targeting to yield EGFP-CaaX or 3xmScarlet-CaaX. For microtubules, EMTB-3xGFP from addgene #26741 was cloned into pBabr, a ΨC31 site-directed transformation vector, between the maternal tubulin promoter and the spaghetti-squash 3′ UTR [35]. These constructs were then integrated into the fly genome at attP2 or attP40 by Rainbow Transgenics Flies, USA, or WellGenetics, Taiwan.

### SILAC metabolic labelling

SILAC metabolic labelling was performed using yeast transformed to produce heavy lysine (SILAC yeast with LysC13/6, Silantes: https://www.silantes.com/). To standardise each of the phosphoproteomic runs for each condition we labelled the proteome of *w^1118^* control flies. Therefore, our analyses included a SILAC-labelled and an unlabelled *w^1118^* extract. To maximise the incorporation of LysC13/6 into the *Drosophila* proteome, unlabelled *w^1118^* adult flies were raised in bottles prepared with SILAC yeast as the only source for amino acids (2% SILAC yeast). The fly media were prepared in agreement with the recommendations of Silantes (www.silantes.com). All emerging larvae were fed on SILAC fly medium from L1 until adult stage, in a temperature (25°C), light and humidity-controlled incubator (Sanyo). The emerging labelled *w^1118^* adults were then transferred to cages for embryo collections, and fed with wet SILAC yeast until disposal of flies (after 2-3 weeks). SILAC *w^1118^* embryos were collected as described above.

### Proteomic analyses

#### Protein digestion

Embryos were lysed in 6M urea and 2M thio-urea (100mM HEPES pH=8.5). Lysates were treated by ultrasonic (20 s, 1 s pulse, 100% power) on ice and cleared by centrifugation (15 min, 22°C, 12.500 x g). Protein concentration was determined with a DC Protein Assay (BioRad).

For each proteome analyses, 200 µg of samples was utilised. Proteins were reduced by Dithiothreitol (22°C, 40 min) followed by protein alkylation using iodacetamide (22°C, 40min in the dark). Lys-C endopeptidase was added for 2h at 22°C. The samples were then diluted to 2M urea using 50 mM ammonium bicarbonate. Trypsin was added in a 1 to 100 enzyme:substrate ratio and incubated overnight at 20°C. Digestion was stopped by acidification using TFA at a final concentration of 0.5%. The resulting peptides were desalted using Waters SPE Columns (C18 material, 50 mg). Peptides were eluted with 60% acetonitrile and 0.1% formic acid. The eluate was dried using a SpeedVac concentrator (Eppendorf) to complete dryness. Peptides were then separated by offline high-pH fractionation.

For phosphopeptide enrichment a SILAC-based quantification was applied. For each sample, 500 ug of protein lysate was mixed with an equal amount of Lys-6 SILAC labelled protein lysate and digested as described above except that Lys-C instead of trypsin was used exclusively. The peptide solution was desalted using Waters SEP-PAK 50 mg C18 cartridges and then subjected for phosphopeptide enrichment.

#### High-pH HPLC offline fractionation

The instrumentation consisted of an Agilent Technologies 1260 Infinity II system including pumps (G7112B), UV detector (G7114A), and a fraction collector (G1364F). A binary buffer system consisting of buffer A, (10 mM ammonium hydroxide in 10% methanol and buffer B (10 mM Ammonium hydroxide in 90% acetonitrile) was utilised. Peptides (resuspended in buffer A) were separated on a KINETEX EVO C18 2×150mm column using a flow rate of 250 µL/min and a total gradient time of 65 min. The content of buffer B was linearly raised from 2% to 25% within 55 min followed by a washing step at 85% buffer B for 5 min. Fractions were collected every 60s in a 96 well plate. Before each run, the system was equilibrated to 100% buffer A. The fractions were then concentrated in a SppedVac concentrator (Eppendorf) and subjected to an additional desalting step using the StageTip technique (SDB-RP, Affinisep). Prior to LC-MS/MS measurement, peptides were solubilized in 10 µL of 2% formic acid and 2% acetonitrile. 3 µL were injected per LC-MS/MS run.

#### Phosphopeptide Enrichment

For phosphopeptide enrichment, the High-Select™ TiO2 Phosphopeptide Enrichment Kit (#A32993) was utilised following the manufacturer’s instructions. In brief, desalted peptides were dried to complete dryness and resuspended in Binding Buffer (included in kit). The peptide solution was centrifuged (10 min, 12.500 x g, 22°C) and the supernatant was transferred to TiO2 tips. Phosphopeptides were enriched and eluted using the provided elution buffer. The eluate was immediately dried in a SpeedVac concentrator (Eppendorf) and stored at −20°C. Prior to LC-MS/MS measurement, peptides were solubilized in 10 µL of 2% formic acid and 2% acetonitrile. 3 µL were injected per LC-MS/MS run.

#### Liquid chromatography and mass spectrometry

The LC-MS/MS instrumentation consisted of a nano-LC 1000 coupled to a QExactive Plus or of a nano-LC 1200 (Thermo Fisher) coupled to a QExactive HF-x instrument via electrospray ionisation. The buffer system consisted of 0.1% formic acid in water (buffer A) and 0.1% formic acid in 80% acetonitrile. The column (75 µm inner diameter, 360 µm outer diameter) was packed with PoroShell C18 2.7 µm diameter beads. The column temperature was controlled to 50°C using a custom-built oven. Throughout all measurements, MS1 spectra were acquired at a resolution of 60000 at 200 m/z and a maximum injection time of 20 ms was allowed. For whole proteome measurements, the mass spectrometer operated in a data-dependent acquisition mode using the Top10 (QExactive Plus) or Top22 (QExactive HF-x) most intense peaks. The MS/MS resolution was set to 17.500 (QE-Plus) or 15.000 (QE-HFx) and the maximum injection time was set to 60ms or 22ms respectively. For phosphoproteome analysis, the MS2 resolutions were set to 30.000 (QEx-Plus) or 45.000 (QEx-Hfx).

#### Proteomic and phosphoproteomic data analysis

Raw files were processed using MaxQuant (v. 1.5.3.8) [61] and the implemented Andromeda search engine [62]. The Uniprot reference proteome for *Drosophila melanogaster* (downloaded: 07.2016, 44761 entries) was utilised. Phosphoproteome and proteome data were analysed separately and the match-between-runs algorithm was enabled. For proteome analysis, the label-free quantification method (MaxLFQ) was enabled using the default settings. For SILAC-based phosphopeptide quantification, a minimum ratio count of 2 was required; the minimum score for the modified peptide was set to 20.

#### Matching and correlation between proteome and phosphoproteome

Protein log2 fold change ratios were matched to the phospho-site table using the Uniprot identifiers. If the phosphorylation site was part of multiple protein groups, the average log2 fold change was utilised. The analysis of the correlation between the fold changes of phosphosites and their host proteins was performed as follows: for each proteome-matched phosphosite, a protein-phosphosite pair was assembled and placed in a scatter plot with 4 quadrants that were connected to 3 possible behaviours: correlation (fold change of protein and phosphosite are both positive (green) or negative (blue), beyond a threshold), anti-correlation (fold change of protein and phosphosite have different signs (red and magenta), beyond a threshold) or no-correlation, fold change of protein resides within threshold range but phosphosite trespasses it (black and white). Finally, we counted the number of protein-phosphosite pairs that displayed each of these behaviours. The threshold used to determine a fold change fluctuation was +/- 0.5 for protein groups, +/- 0.35 for phosphosites (see below Hierarchical clustering analyses, Threshold determination).

### Immunostainings and live imaging procedures

#### Synchronised egg collections

Eggs were collected for 1 h, allowed to develop for a further 2hs 30’ in a temperature (25°C), light and humidity-controlled incubator (Sanyo) and then dechorionated in sodium hypochlorite (50% standard bleach in water) and washed thoroughly with water. Depending on the type of staining and antigen, embryos were fixed using the appropriate standard protocols.

#### In situ RNA hybridisation

Antisense probes for Dpp, Sog and Snail were used on dechorionated embryos by applying *Drosophila* standard protocols for *in situ* hybridisation with digoxigenin-labelled RNA-probes [63].

#### Heat fixation for imaging of Armadillo/β-Catenin

Dechorionated embryos were transferred to a beaker containing 10ml of boiling heat-fixation buffer (For 1L in water: 10X Triton-Salt Solution, 40g NaCl, 3ml Triton X-100 (T9284 Sigma)), and fixed for 10 sec. Fixation was stopped by placing the beaker containing the embryos on ice.Vitelline membranes were removed by transferring the embryos to a tube containing heptane:methanol (1:1), vortexed for 30 sec. and rehydrated.

#### Fixation for imaging of microtubules

To visualise microtubules, a formaldehyde-methanol sequential fixation was performed as previously described [50]. Dechorionated embryos were fixed in 10% formaldehyde (methanol free, 18814 Polysciences Inc.) in PBS:Heptane (1:1) for 20 min at room temperature (RT), and devitellinised for 45 sec in 1:1 ice-cold methanol:heptane. Embryos were stored for 24 hs at −20°C and rehydrated before use.

#### Antibody staining procedures

Rehydrated embryos were blocked for 2 hs in 2% BSA (B9000, NEB) in PBS with 0.3% Triton X-100 (T9284 Sigma). Primary antibody incubations were done overnight at 4°C. Primary antibodies used were: mouse anti α-tubulin 1:1000 (T6199, clone 6-11B-1, Sigma), mouse anti acetylated-α-tubulin FITC conjugated 1:250 (sc23950, Santa Cruz Biotechnology), rat anti tyrosinated-α-tubulin 1:250 (MAB1864-I, clone YL1/2, EMD Millipore/Merck), mouse anti-Armadillo/β-Catenin 1:50 (N27A1, Developmental Studies Hybridoma Bank), rabbit anti Snail [64] 1:500. Incubations with secondary antibodies were performed for 2 hours at RT. Alexa Fluor 488- and 594-coupled secondary antibodies were used at 1:600 (488 and 594 Abcam).

#### Preparation of physical cross-sections

Immunostained embryos embedded in Fluoromount G (SouthernBiotech 0100-01) were visually inspected under a dissecting microscope (Zeiss binocular) to select the desired developmental stages. The embryos were sectioned manually with a 27G injection needle at approximately 50% embryo length and slices were mounted for microscopy.

#### Image acquisition

Images in Figure 1C were acquired with a Zeiss LSM880 Airyscan microscope, using a Plan-Apochromat 63x oil (NA 1.4 DIC M27) objective at 22°C, with a z-slice size = 0.3 μm. Acquired volumes were max-projected (along z axis) for a range of 1.5 μm (5 slices). Images in Figure 7A and Supplementary Fig. 5B were acquired using a super resolution Deltavision OMX 3D-SIM (3D-SIM) V3 BLAZE from Applied Precision (a GE Healthcare company). Deltavision OMX 3D-SIM System V3 BLAZE is equipped with 3 sCMOS cameras, 405, 488 and 592.5 nm diode laser illumination, an Olympus Plan Apo 60X 1.42 numerical aperture (NA) oil objective, and standard excitation and emission filter sets. Imaging of each channel was done sequentially using three angles and five phase shifts of the illumination pattern. The refractive index of the immersion oil (Cargille) was 1.516. Acquired volumes were max-projected (along z axis) for a range of 3 μm (10 slices). Images in Figure 7B and Supplementary Fig. 5A were acquired with a Leica SP8 microscope equipped with white laser. Gated detection on HyD detectors was used for each shown channel using a Plan-Apochromat 63x oil (NA 1.4) objective at 22°C, with a z-slice size = 0.3 μm. Acquired volumes were max-projected (along z axis) for a range of 1.5 μm (5 slices).

#### Live imaging of myosin light chain

Dechorionated embryos expressing maternally provided sqh::mCherry were mounted in 35mm glass-bottom petri dishes in two different ways with either the embryonic dorsal, lateral or ventral surface facing the glass bottom or vertically glued (heptane glue) on their posterior end to the glass-bottom and embedded in 0.8% low melting point agarose in PBS that was previously cooled to 30°C. For the superficial imaging of the sub-apical domain of embryos along their dorsal, lateral and ventral sides, we acquired volumes of 20 μm (z-slice size of 0.8 μm) using a PerkinElmer Ultraview ERS (microscope stand Zeiss Axiovert 200) with a Yokogawa CSU X1 spinning disk with a Plan-Apochromate 63x (NA 1.4, oil) objective at 22°C. The vertical mounting enabled the imaging of myosin light chain along the dorso-ventral cross-section of living embryos [65] in the x-y plane. We acquired 1-3 slices (z-slice size = 1,1um) at 130-150 μm from the posterior end of the embryo using a Zeiss LSM780 NLO 2-photon microscope with a Plan-Apochromat 63x objective (NA 1.4, oil, DIC M27) at 22°C.

### Calculation of correlation matrices

Correlation matrices were calculated using Matlab R2019b on the proteins and phosphosites that were detected in all genotypes and all the replicates. For this, we used the ‘corr’ function to calculate the Pearson correlation between log2 intensities of proteins or phosphosites in the replicates of all genotypes. The resulting correlation matrix was plotted using the ‘clustergram’ function by applying the following settings: linkage: average; RowPDist & ColumnPDist: correlation.

### Development of a linear model for protein and phosphosite abundances

We defined a ‘linear model’ that is based on the assumptions that the three types of mutant embryo each represent one region along the DV axis of the embryo, and that protein abundance in the three regions should add up to the total protein abundance in the entire embryo, and therefore, the abundance in the three mutants should add up to the same value, if each is weighted by the region occupied in the wildtype embryo.

This linear model can be expressed for each protein ProtX as a sum, ^t-^wt_Prot X_, where D, L and V are the abundance of ProtX in the proteomes of the three mutant genotypes (means of the Log2 intensity values of the replicates, transformed to its linear value), and a, b and c represent the proportion of each region along the dorsoventral axis:.

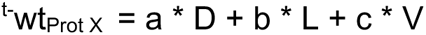

This model requires values a, b, and c for the three regions along the DV axis. For the mesoderm, this has been reported as 0.2 from measurements on cross-sections [21], but we wanted to determine the theoretical optimum for each of the values. The theoretical optimum would be one for which the proportions for the three regions when used in the sum yield a theoretical value ^t^wt_Prot X_ that is closest to the experimentally measured value ^m^wt_Prot X_ for that protein in the wildtype embryo.

To systematically explore the proportions for each region, we tested all possible combinations for a, b and c at 0.05 steps in the range from 0 to 1 (i.e.: 0, 0.05, (…), 0.95,1). For each calculated ^t-^wt_Prot X_, we calculated the deviation from the ideal by subtracting the mean linear intensity measured for the same protein in a wild type embryo (^m^wt_Prot X_), and transforming this difference between the theoretical and measured wild type to Log2 scale as follows:

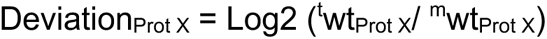

For each possible combination of a, b, and c we obtained distributions of Deviation_Prot X_ values, for which we calculated the Interquartile Range (IQR) using the IQR function from Matlab 2019b. Based on the assumption that the best matching proportions should lead to the narrowest dispersion of the distribution of Deviation_Prot X_, we sorted the combinations of proportion constants based on their calculated IQRs. The combination with the smallest IQR and a ventral domain (‘c’) proportion constant of 0.2 was selected for the linear model.

Because we used two different ventralising genotypes (*Toll^10B^* and *spn27A^ex^*), the ^t-^ wt_Prot X_ was calculated twice for each protein, one for for each mutant genotype combination (i.e.: D-L-V*_Toll10B_* and D-L-V*_spn27A_*).

### Hierarchical clustering analyses

#### Data generation and hierarchical clustering

We included in this analysis all proteins that were detectable in the wildtype (5883/6111), even if they were undetectable in one or more mutant populations. To obtain clusters that represented the behaviours of proteins and phosphosites with respect to the wild type genotype, we took for each protein and phosphosite the mean Log2 fold change. For this, we first calculated the mean Log2 intensity values per protein and genotype, and next, calculated the Log2 fold changes (FCs) for each protein and phosphosite between each DV mutant genotype and the wild type. Because we wanted to cluster exclusively by the direction but not the extent of changes between the DV mutant genotypes and the wild type, we assigned to each FC a value (1, −1 or 0) based on exceeding a FC threshold (see below ‘*Threshold determination*’). For proteins, the threshold was +/- 1.4 (0.5 in Log2) FC, for phosphosites was +/- 1.27 (0.35 in Log2) FC. When proteins and phosphosites exceeded the FC threshold, we assigned a +1 or −1 for positive and negative FCs respectively. When proteins and phosphosites remained within the threshold range (i.e.: for proteins: −1.4 (0.5) < FC < 1.4 (0.5); for phosphosites: −1.27 (0.35) < FC < 1.27 (0.35)), we assigned a 0. For proteins and phosphosites that were undetected in a particular DV patterning mutant, we assumed -based on the detection of Twist and Snail across mutant genotypes-, these were in decreased abundance vs. the wild type, and we assigned to these −1.

Next, we filtered the set of proteins and phosphosites that we used as a source for the hierarchical clustering. Proteins and phosphosites with a 0 assigned in all FC comparisons (i.e. D vs. WT = 0; L vs. WT = 0, Vtl vs WT = 0, Vsp vs WT = 0) were considered unchanged in our study and therefore were filtered out (N° proteins = 2156/6111; N° phosphosites = 1234/6259). Proteins and phosphosites with a +1 or a −1 assigned to all FC comparisons (i.e.: D vs. WT =1 (−1); L vs. WT = 1 (−1), Vtl vs WT = 1 (−1), Vsp vs WT = 1 (−1)) were also filtered out (N° proteins = 329/6111; N° phosphosites = 615/6259). We proceeded with the clustering of the rest of proteins (3398/6111) and phosphosites (3433/6259).

The thresholded FCs of the filtered set of proteins and phosphosites were transformed in row zscores (i.e.: calculated per protein and per phosphosite). The hierarchical clustering was conducted both on rows (proteins or phosphosites) and columns (FC of each mutant genotype vs. the wildtype) using the ‘clustergram’ function (Matlab R2019b) setting the linkage parameter in ‘average’ and the row probability distance parameter in ‘Euclidean’. The output of the clustering in the figures was set to be displayed using the mean Log2 FCs of proteins and phosphosites between the DV mutant genotypes and the wild type.

#### Threshold determination

We determined the threshold to be applied to the FCs between the DV mutant genotypes and the wild type of each proteomic experiment by analysing the variability of the FCs within each biological replicate. We first took the mean Log2 intensity per protein for the wildtype ‘^WT^Mean_ProtX_’, and next, calculated the Log2 FC between the Log2 intensity of each technical replicate with measurable intensity for each biological replicate and the ^WT^Mean_ProtX_ as follows:

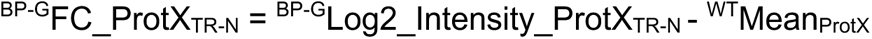

where BP-G is the biological replicate of a particular genotype and TR-N a technical replicate (1 < N < 3) for a biological replicate BP-G. Therefore, we obtained a set of ^BP-G^FC_ProtX_TR-N_ values per protein, and the number of ^BP-G^FC_ProtX_TR-N_ values per protein equals the number of technical replicates with measurable intensity for a particular biological replicate.

Next, we calculated ^BP-G^stdev_ProtX, which is the standard deviation of the set of ^BP-G^FC_ProtX_TR-N_ values per protein and for each biological replicate. In this way, we obtained a distribution of the standard deviation of FCs per biological replicate. For each biological replicate distribution of FCs standard deviations we calculated the IQR (Prism Graphpad V8) and extracted the 3rd quartile value (^BR-G^Q3) to capture up to 75% of the variability of each distribution:

| Biological Replicate Proteome | 3rd Quartile Value (Q3) |
| --- | --- |
| Dorsalized (replicates 1-3) | 0.543 |
| Dorsalized (replicates 4-6) | 0.492 |
| Lateralized | 0.471 |
| Ventralized *spn27A* (replicates 1-3) | 0.463 |
| Ventralized *spn27A* (replicates 1-6) | 0.481 |
| Ventralized *Toll*<sup>10B</sup> | 0.565 |

| Biological Replicate Phosphoproteome | 3rd Quartile Value (Q3) |
| --- | --- |
| Dorsalized | 0.405 |
| Lateralized | 0.366 |
| Ventralized *spn27A* | 0.377 |
| Ventralized *Toll*<sup>10B</sup> | 0.259 |

Finally, we defined the FC threshold as the mean of the Q3 value across the biological replicates of each experiment (number of: biological replicates proteome = 6; biological replicates phosphoproteome = 4) as follows:

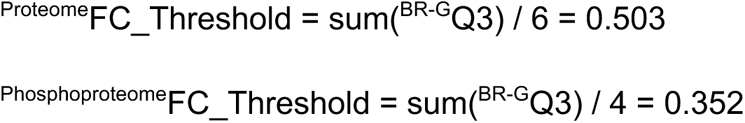

### RNA-proteome comparison along DV cell populations

#### Data generation: NovoSpark analyses of single-cell RNAseq data

The single-cell RNAseq data derived from stage 6 *Drosophila* embryos [29], were spatially reconstructed with novoSpaRc [66]. As prior spatial information 84 known gene expression patterns were used from the BDTNP atlas (downloaded from: https://shiny.mdc-berlin.de/DVEX/). NovoSpaRc embeds each cell probabilistically over 3039 locations using a generalized optimal-transport approach. This results in a ‘RNA Atlas’, which includes a predicted spatial gene expression pattern for every detected gene.

#### RNA clustering

We excluded all genes that were scored as ubiquitously expressed in THE RNA ATLAS [29]. Of the remaining 8924 genes, we selected those that were also listed in our clustered proteomic dataset (3346 genes coding for 3383 proteins), yielding a list of 3086 non-ubiquitous genes that were present both in the RNA atlas and the clustered proteome. These sorted into classes by comparing the expression pattern of each to that of six reference genes with restricted dorso-ventral expression that represent the six regulation categories D, L, V, DL, DV and VL. We used as reference genes *dpp* for dorsal, the average between *sog* and *soxN* for lateral, *twist* for ventral, *crb* for dorsal+lateral, *net* for dorsal+ventral and *neur* for lateral+ventral. To compare similarity for each of the 3086 genes we calculated their spatial Pearson correlation to each of the reference genes. Each gene was then classified as belonging to the category of the reference gene for which the Pearson correlation was the highest. We therefore obtained 6 clusters of genes, which we termed ‘DV RNA clusters’ each of them with their corresponding maximum Pearson spatial correlation value. To filter out false positives, we selected those genes with the largest similarity to the reference genes, for which we expected strong differential expression along the dorso-ventral axis. We did this by determining the Pearson spatial correlation value corresponding to the 95th Percentile of each dorso-ventral RNA cluster, using the ‘prctile’ function (Matlab R2019b). Finally, we used the 95th percentile value as a threshold to filter the 5th percentile of genes from each DV RNA cluster. We obtained a list of 155 genes that we used to compare against the proteome clusters (Supplementary Table 8), which we termed ‘DV RNA Reference Sets’.

#### RNA-proteome comparison

The filtered set of 155 genes codes for 157 proteins. We grouped the 157 proteins based on their DV cluster assignment and for each DV cluster, we classified its proteins based on the RNA reference set to which their genes had been allocated. For DV clusters 1-12, we classified the results of this comparison as ‘perfect match if the RNA expression pattern and the DV cluster belong to the same regulation category, as ‘partial match if the RNA expression pattern and the DV cluster coincided only in one DV domain, or as mismatch if the RNA expression pattern and the DV cluster belong to mutually exclusive regulation categories.

### Calculation of euclidean distance score

We developed a score based on a calculated Euclidean distance to measure the proximity of each protein in a particular DV cluster to the most extreme fold changes measured in DV mutant vs. wild type comparisons. We used the same approach for the phosphosites.

We first calculated the mean log2 fold change (FC) between the DV mutants and the wild type (which meant we could not assess proteins nor phosphosites that were not detected in the wild type). Next, we rescaled each of the FC distributions [dorsalized (D) vs. WT, lateralized (L) vs. WT, ventralized *Toll^10B^*/def (Vtl) vs. WT and ventralized *spn27A^ex^*/def (Vsp) vs. WT] to transform the log2 FC = 0 in the original distributions as log2FC = 0.5 in the new, rescaled distribution. We first identified the upper and lower limits of each FC distribution, and next, transformed each pair (upper and lower) of limits to their absolute values. This enabled the identification of the largest absolute limit for each FC distribution, and depending whether the upper or the lower limit was the absolute largest, we used one of the following equations to rescale:

If the upper limit of a particular FC distribution is the absolute largest:

i. log2FC_rescaled_i = (log2FC_original_i + |max_log2FC_original|) / (2* |max_log2FC_original|)

If the lower limit of a particular FC distribution is the absolute largest:

ii. log2FC_rescaled_i = (log2FC_original_i + |min_log2FC_original|) / (2* |min_log2FC_original|)

Where ‘i’ is a particular protein, and max_log2FC_original and min_log2FC_original are the upper and lower limit values for a particular FC distribution. Using this rescaling approach, we obtained a new set of rescaled FCs for each distribution (D vs. WT, L vs. WT, Vtl vs. WT and Vsp vs. WT). We considered those proteins that were undetected in a mutant genotype as being in decreased abundance in that genotype, and imputed the lower limit value of the rescaled FC distribution for that genotype.

For each protein, we assigned two vectors with their corresponding rescaled FCs (shown here for one ventralized genotype, but also calculate separately for the other):

(D_rescaledFC_i, L_rescaledFC_i, Vtl_rescaledFC_i)

Next, we assembled reference vectors representing the most extreme behaviours for each regulation category (Fig. 3D, upper panel), using the rescaled FCs:

| Regulation category | DV cluster | Reference vector components | Reference vector values with Vtl* | Reference vector values with Vsp* |
| --- | --- | --- | --- | --- |
| Dorsal | 1 | (max<sup>rFC</sup>, min<sup>rFC</sup>, min<sup>rFC</sup>) | (1, 0, 0.0047) | (1, 0, 0.0485) |
| Lateral | 2 | (min<sup>rFC</sup>, max<sup>rFC</sup>, min<sup>rFC</sup>) | (0.1127, 0.9865, 0.0047) | (0.1127, 0.9865, 0.0485) |
| Ventral | 3, 4, 5 | (min<sup>rFC</sup>, min<sup>rFC</sup>, max<sup>rFC</sup>) | (0.1127, 0, 1) | (0.1127, 0, 1) |
| Dorsal+Lateral | 6 | (max<sup>rFC</sup>, max<sup>rFC</sup>, min<sup>rFC</sup>) | (1, 0.9865, 0.0047) | (1, 0.9865, 0.0485) |
| Dorsal+Ventral | 7, 8, 9 | (max<sup>rFC</sup>, min<sup>rFC</sup>, max<sup>rFC</sup>) | (1, 0, 1) | (1, 0, 1) |
| Lateral+Ventral | 10, 11, 12 | (min<sup>rFC</sup>, max<sup>rFC</sup>, max<sup>rFC</sup>) | (0.1127, 0.9865, 1) | (0.1127, 0.9865, 1) |

Where max^rFC^ and min^rFC^ are the maximum and minimum rescaled FCs for the corresponding distributions (D vs. WT, L vs. WT, Vtl/Vsp vs. WT). Finally, we calculated the Euclidean distance score (ED Score) between each protein in clusters 1 to 12, and the reference vectors that correspond to each DV class:

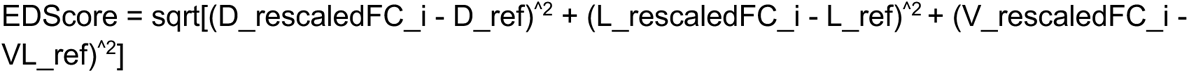

Where D_ref, L_ref and V_ref are the reference vector values* for the corresponding regulation category.

### Ontology analyses using diffused networks

#### Data generation

The method employed here is similar to the one developed in Giudice et al [32]. Briefly, we retrieved the *Drosophila* protein-protein interaction network from IntAct (last update June 2020). We modelled the edge weights using the Resnik semantic similarity is employed to model the edge weights [67] semantic similarity calculated employing using the Semantic Measures Library [68]. We also generated 1000 random networks, where the node degrees are conserved, employing the vl method from the igraph library [69]. The edge weights of the random network are updated accordingly. To correct for the hub bias we applied the Laplacian normalisation to all networks using:

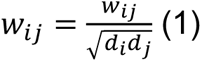

Where *w_ij_* indicates the edge weight and *d_i_* and *d_j_* represent the weighted degree of node *i* and node *j* respectively. We extracted from the Pfam [70] database (last update June 2021), all the kinases detected in *Drosophila* by selecting the CL0016 clan. Next, we employed the UniprotKB database to distinguish the tyrosine kinases (family: PF07714) from other kinases. In total 56 tyrosine kinases and 251 other kinases are present in the network. We also precalculated the mean and the standard deviation of the Resnik semantic similarity of each regulated node in the network against each other.

#### Seed selection and network diffusion

We applied the random-walk-with-restart-based algorithm [32] for the following DV clusters: D (1), L (2), V (5), DL (6), DV (9) and LV (12), once each for the ED score and once for the deviation values, and each for the protein and phosphoproteomic datasets. In the case of the deviation values, we used only those proteins or phosphosites within the interquartile range. We assigned as seed value the reciprocal of the absolute value from both the ED Score or the Log2 Fold Change Deviations. Note that if multiple phosphosites are assigned to the same protein we selected the median of the ED scores or Log2 Deviations. We then partition the seed set in tyrosine kinases, remaining kinases and other proteins and perform the random walk with restart (RWR) from each of the three partitions separately. We also repeated the same procedure with the same set of initial nodes against 1000 random networks. We estimated the empirical p-value for each node of the network as the percent of its random scores that exceed the real score and selected only the nodes with a p-value<0.05 in at least one of the three partitions. The resulting subnetworks are further filtered using the ego decomposition [32]. Briefly, for each seed node we extracted ego networks with a maximum distance of 2 steps from the ego. We then filtered the ego networks by selecting only the most similar functional nodes to the ego using this formula:

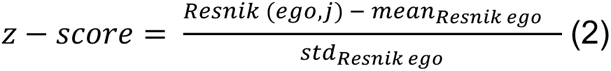

Where *Resnik*(*ego*, *j*) is the semantic similarity between the ego and a node j in the ego network, *mean_Resnik ego_ and std_Resnik ego_* are the mean and the standard deviation of the ego against all the other nodes in the initial network. Nodes with a z-score>1.28 (equivalent to a pvalue<0.1) are retained. After this filtering, the resulting ego networks with less than 5 nodes are discarded. Additionally, the weight of the ego networks are changed according to (3) to reflect the functional impact of the dysregulation of the ego on the neighbouring nodes.

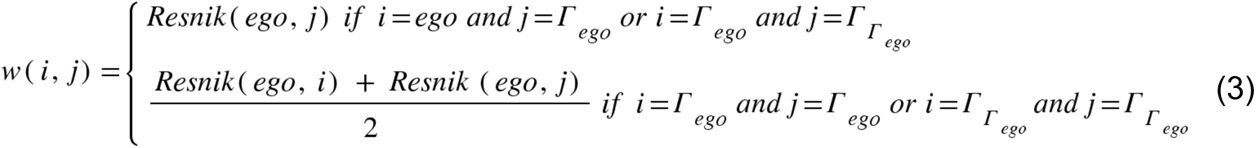

where Γ(ego) represents the nodes at distance 1 from the ego and Γ_Γ(ego)_ represents the nodes at distance 2 from the ego. The ego networks obtained are normalised again to correct for hubs using the Laplacian normalisation using (1). For each ego network, we then calculate the topological distance vector and the functional distance vector as in Giudice et al. The topological distance vector is calculated using the following formula (4):

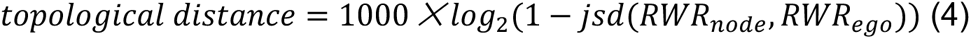

Where *jsd* refers to the Jensen-Shannon distance, representing the similarity between two probability distributions. The RWR_node_ refers to the RWR probability vector when one of the nodes of the ego network is selected as seed, and the RWR_ego_ refers to the RWR probability vector when the ego is the seed node. The functional vector is defined as the logarithm of the semantic similarity between the ego and any other nodes in the network (5).

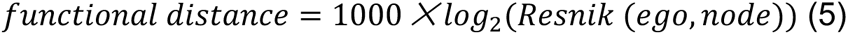

Where *Resnik(ego, node)* represents the semantic similarity measure between the ego and the node under consideration. To assess the most similar nodes to the ego, the Kernel Density Estimation (KDE) measure (with Gaussian kernel and bandwidth estimated using the Silvermann formula) to assess the most similar nodes to the ego, is employed. KDE estimates the probability density function (PDF) of the topological and semantic similarity vectors obtained at the previous step. For each ego network we selected only those nodes within a 0.7≤PDF≤1.0 of both topological and functional similarity. All the nodes overcoming this threshold are selected for the enrichment analysis against the cellular component domain of GO.

### Ontology analyses of extreme deviating proteins and phosphoproteins using PANTHER protein class

We filtered the proteins and phosphosites with an absolute Log2 deviation value larger than the 95^th^ percentile (prctitle Matlab function) of the distribution of absolute log2 deviation values. Because deviation values were calculated separately for each ventralized genotype (*Toll^10B^*/def or *spn27A^ex^*/def), we obtained two independent lists of proteins and phosphosites with deviation values exceeding the 95^th^ percentile. From these lists, we selected the proteins and phosphosites whose log2 deviations exceeded the 95^th^ percentile threshold with both ventralized genotypes. For phosphosites, we used the host phosphoprotein for the ontology analyses. When two or more phosphosites with large deviations were hosted by the same phosphoprotein, we counted the phosphoprotein only once. We therefore obtained 206 proteins and 154 phosphoproteins (191 phosphosites) that were used in the ontology analyses.

The ontology analyses were performed using the PANTHER platform (http://www.pantherdb.org/, release PANTHER 17.0 dated February 23^rd^ 2022). We queried the ‘Functional classification gene list’, based on ‘*Drosophila melanogaster*’ organism data. We used the FBgn (Flybase Gene Number) of the proteins and phosphoproteins to produce the query in Panther, and focused on the ‘Protein Class’ classification. The protein class query allocated 125/206 proteins and 110/154 phosphoproteins to protein class terms. Using gene ontologies from Flybase we manually allocated 48/206 proteins and 35/154 phosphoproteins to one or more of the 24 parental protein class categories. 33/206 proteins and 9/154 phosphoproteins could not be allocated to any parental class category, remained unassigned and were therefore excluded from the reported analyses. In summary, the reported protein class ontology analyses of extremely deviating proteins and phosphoproteins is based on 173/206 proteins and 145/154 phosphoproteins.

### Functional perturbations on microtubules

#### Depolymerisation of microtubules and imaging

Embryos laid by flies heterozygous for EMTB-3xGFP and 3xmScarlet-CaaX transgenes or EGFP-CaaX and H2Av-mRFP1 [71] transgenes were submerged under Halocarbon oil 27 (Sigma-Aldrich) for staging. Early cellularizing embryos were selected, dechorionated with 50% bleach after removal of Halocarbon oil, washed with H_2_O, mounted with heptane glue on a coverslip, desiccated with silica gel or Drierite for 10’-15’, and subsequently covered with a 3:1 mixture of Halocarbon 700 and 27 oils. For Colcemid injection, 4 mg/ml Demecolcine (Sigma-Aldrich) in H_2_O was injected with a custom-made injection needle that was prepared from a borosilicate glass micropipette (Drummond) with a Sutter Instrument pipette puller (P-97/IVF) and a Narishige grinder (EG-44). The stage of injection was controlled based on the transmission brightfield image. A volume of ∼65 pL, measured with a 20X dry lens via an objective micrometre, was injected into the middle section of embryos. Injection was performed with a Narishige IM400 setup mounted on a Nikon Ti2/Eclipse inverted microscope equipped and under a 60x/NA1.42 oil immersion objective. Imaging was performed on a Yokogawa CSU-W1/SORA imaging system mounted on the same scope. Two laser lines (488 and 561 nm) were used to excite the sample, while a tandem of sCMOS cameras (Prime BSI, Teledyne Photometrics) were used to acquire the image with 2×2 binning. A single z-stack volume was acquired prior to injection, followed by a post-injection z-stack time series. The gap between pre-injection and post-injection imaging was typically 2’∼3’. CSU-SORA 4x zoom was used for imaging EMTB-3xGFP and 3xmScarlet-CaaX with a z-step size of 0.3 μm and a total z-depth of 6.3 μm at a rate of 30s per volume, while CSU-W1 was used for EGFP-CaaX and H2Av-mRFP1 with a z-step size of 1 μm and a total z-depth of 40 μm at a rate of 1’ per volume.

#### Image processing and quantitation

For quantitation of nuclear position, single z-slice H2Av-mRFP1 images were converted into tif format, blurred with a Gaussian filter (σ=2), and segmented with CellPose (v2.2) in 2D using a custom-pretrained nuclear model tailored for each side of the embryo based on manual correction on segmentation generated by a default nuclei model with nuclear diameter set as 20 pixels. The stitch mode was used with a stitch threshold of 0.4 to generate 3D nuclear segments. Nuclear segments were filtered by size (1000-5000 pixels) and height (>10 μm), while those located at the edge of the imaging area were excluded for data processing. The regionprops function implemented in the Skimage Python library was used to define the bounding box of each nuclear segment, from which the middle Z coordinate of the bounding box was designated as the nuclear position. For time alignment, t_0_ (the onset of gastrulation) was defined as the time point, at which apical constriction in ventral furrow produces a 2.5 μm gap between the cell apex and vitelline membrane for the datasets acquired on the ventral side. Using this t_0_ designation (from water-injected embryos imaged on the ventral side), the pre-injection cellularisation depths were fitted to a linear function based on the assumption that cellularisation depth is linear with time during mid-cellularisation. For datasets acquired on the lateral and dorsal sides, the pre-injection timing relative to the onset of gastrulation was derived by plugging in the pre-injection cellularisation depth, from which the t_0_ frame of the dataset was derived. Data processing and plotting were performed with custom-made Python codes using Numpy, Pandas, Matplotlib, and Seaborn libraries.

For *en face* membrane visualisation, 3xmScarlet-CaaX images were deconvolved using the Huygens Software (Scientific Volume Imaging) with the deconvolution algorithm Classic MLE using custom parameter sets.

### Data availability

Supplementary tables information are available upon request in Zenodo public repository under DOI: ‘//doi.org/10.5281/zenodo.8278666’.

---

**Supplementary Figure 1. Proteomic and Phosphoproteomic strategy in *Drosophila* embryos at the point of gastrulation. A)** Images (confocal, max-projected) of physical cross-sections of heat-fixed embryos showing the transition from stage 5 (late cellularisation) to stage 6, and the progression of gastrulation during stage 6 (ventral furrow formation), stained using antibodies against β-Catenin/Armadillo (green in top panel; greyscale in bottom panel) and Snail (magenta in top panel).

**B)** Scheme explaining the strategy for proteomic and phosphoproteomic experiments. Synchronised collections of embryos from wild type or mutant mothers, representing the dorsal ectodermal (dorsalized: *gd^9^*), neuroectodermal (lateralized: *Tl^rm9^*/ *Tl^rm10^*) and mesodermal (ventralized: *Toll^10B^*/*def* & *spn27A^ex^*/*def*) cell populations were allowed to develop for 2hs 30’ at 25°C, manually selected under visual inspection for 15’ to secure the collection of stage 6 embryos and immediately frozen in liquid nitrogen. For the phosphoproteomic experiments, we used SILAC metabolic labeling to optimise the quantification strategy.

**C)** Workflow for quantitative label free proteomics using high-pH offline peptide fractionation followed by LC-MS/MS of individual fractions.

**D)** Workflow for quantitative SILAC-phosphoproteomic analyses. Phosphopeptide enrichment was performed using TiO2 beads, and LC-MS/MS based identification and quantification. For the mass-spectrometry analysis of each replicate, we combined equal amounts of protein from wild type embryos labeled with SILAC-Lys 13/6, and embryos of the target genotype (500 μg target genotype : 500 μg SILAC wild type).

**E)** Histogram showing the distribution of the number of protein groups with respect to the log2 fold change between the intensity values measured for a given protein group in SILAC wild type vs. non-SILAC wild type embryos. H stands for heavy (or SILAC) and L stands for light (or non-SILAC) in pilot runs.

**Supplementary Figure 2. Proteomic validation of dorso-ventral embryonic cell populations**

**A)** Log2 intensity (top) and log2 fold change (FC, bottom) of Toll and Spn27A proteins.

**B)** Log2 intensity (top) and log2 fold change (bottom) of Toll phosphosite S871.

**C)** Log2 intensity (top) and log2 fold change (bottom) of additional mesodermal proteins.

**D)** Log2 intensity (top) and log2 fold change (bottom) of additional ectodermal fate determinants.

**E)** Log2 intensity (top) and log2 fold change (bottom) of Cactus protein (left panels) and Cactus phosphosites (right panels): S463 (yellow dots), S467 (green dots) and S468 (magenta dots).

Colors depict DV mutant genotypes and their corresponding comparisons against wild type: blue: dorsalized, magenta: lateralized, yellow: ventralized (*Tl^10B^*/def and *spn27A^ex^*/def). Bars depict mean and standard error of the mean across replicates. Absence of a dot indicates the protein was not detected in a particular condition or log2 FC calculation not feasible, absence of error bars in Log2 intensity indicate protein was detected in a single replicate.

Dotted line indicates log2 FC = 0, log2 FC = 0.5 (for proteins) or log2 FC = 0.35 (for phosphosites).

**Supplementary Figure 3. Proteomic validation of dorso-ventral embryonic cell populations A)**. The protein-level expression of the transcription factor Snail in DV mutants phenocopies the expression pattern across DV cell populations in wild type embryos. Images (confocal, max-projected) of physical cross-sections from heat-fixed embryos showing Snail (grayscale) antibody signal in wild type, dorsalized (D), lateralized (L) and ventralized embryos (Vtl: *Tl^10B^*/def and Vsp: *spn27A^ex^*/def). **B and C)** Outcome of the systematic exploration for the optimal combination of cross-sectional proportions for dorsal (D), lateral (L) and ventral (V) cell populations, with each population being varied in steps of 0.05. On the x axis, the 233 possible combinations are plotted in the order of the proportions assigned to the D population, and the remaining proportion distributed between L and V. Only the values for D are indicated on the axis, see Suppl. Table 6 for the L and V values. The y axis shows the Interquartile Range (IQR) of the distribution of the Log2 Deviations obtained with each possible combination of D, L and V proportions. Calculations were performed independently for the *Toll^10B^/def* (B) or *spn27A^ex^/def* (C) data. Light blue dots indicate the best combinations with a ventral domain proportion (V = 0.2) that matches the experimentally determined region occupied by the mesoderm. **D)** Pie charts showing the protein class ontology (http://www.pantherdb.org/) of proteins with the highest absolute deviations (95th percentile). Inset pie chart shows the children ontology terms for the ‘Metabolite interconversion enzyme’ class. **E)** Pie charts showing the protein class (Panther) ontology of the host proteins for phosphosites with the 5% most extreme absolute deviations (> 95th percentile). The left pie chart shows the children ontology terms for the ‘Metabolite interconversion enzyme’ class.

**Supplementary Figure 4. Methodology and supporting data for the comparison between RNA and protein abundance.**

**A)** Methodology for clustering non-ubiquitous genes according to their dorso-ventral RNA expression.

**B)** Swarm plot showing the distribution of the maximum Pearson spatial correlation values, used for the assignment of non-ubiquitous genes to a particular DV RNA sets. Each dot is a gene. DV RNA sets are mutually exclusive. Bars indicate median and the inter-quartile range (IQR).

**C)** Number of non-ubiquitous genes allocated to each DV RNA reference set using the Pearson spatial correlation to reference genes (panel B and Fig. 4A).

**D)** Maximum Pearson correlation values for increasing percentiles within the distributions of each RNA reference set. To assemble the DV RNA reference sets (Fig. 4C), we filtered the genes that at the RNA level displayed the largest variation along the dorso-ventral axis. For this, we selected genes with a maximum Pearson correlation value larger than the 95% percentile (arrow) of the corresponding DV RNA set distribution.

**E)** Matching of genes in the RNA reference sets with the proteome DV clusters in which the ventralized mutants display a non-consistent behaviour against the wild type (DV clusters 3, 4, 7, 8, 10 and 11, see Fig. 3B,C). Coloured pie charts represent the allocation of genes within each DV cluster to the RNA reference sets (colour code as in Fig. 3C). Grayscale pie charts represent the outcome of the comparison for filtered genes: white: perfect for an exact match, grey: partial, if the RNA reference included the correct match but also another region, black: mismatch, where the protein expression did not overlap with the RNA reference. Values in the centre of pie charts indicate the number of genes compared.

**F)** Comparison between the identity of genes in the RNA reference sets with DV clusters 13 and 14, (Fig. 3B,C), with proteins that displayed increased abundance in all DV mutants (vs. wild type) with the exception of either of the ventralized genotypes (DLVtl or DLVsp). Coloured pie charts represent the allocation of genes within these DV clusters to the DV RNA reference sets (Blue: dorsal, magenta: lateral, yellow: ventral, cyan: dorsal+lateral, green: dorsal+ventral, pink: lateral+ventral).

**Supplementary Figure 5. Cellular component terms significantly enriched in diffused networks.** Heatmap representation of the significantly enriched cellular component terms significantly enriched in a single network across all DV clusters. In A) filtered out proteome ontology terms; in B) filtered out phosphoproteome ontology terms.

**Supplementary Figure 6. Microtubule analyses during gastrulation and calibration of colcemid injections.**

**A)** Image (confocal, max-projected) of a physical cross-section from fixed embryos stained with antibodies against tyrosinated (magenta) and acetylated (green) α-tubulin during ventral furrow formation. Arrow: detection of acetylated α-tubulin specifically in basal-lateral microtubules (inverted basket); blue arrowhead: tyrosinated α-tubulin in sub-apical microtubules; red arrowhead: tyrosinated α-tubulin in basal-lateral microtubules. Insets show acetylated and tyrosinated α-tubulin in stretching and constricting mesodermal cells; arrowheads/arrow in insets show differential acetylation and tyrosination of microtubules in a mesodermal cell undergoing stretching. Scale bar is 50 μm.

**B)** Images (OMX super-resolution microscope, max-projected) of stretching mesoderm cells (ventral domain) from fixed embryos stained using antibodies against E-cadherin (magenta in top panels; grayscale in middle panels) and α-Tubulin (green in top panels; grayscale in bottom panels). A stretching mesodermal cell (yellow-coloured) was cropped and shown with superficial (left panel) and resliced (to show ‘y-z’ planes) views. The ventral furrow is located to the right of the field of view. Arrow indicates a microtubule in the sub-apical domain that is bent towards the direction of cell stretching (furrow). Scale bar is 10 μm.

**(C, D)** Time lapse series of microtubules labelled with EMTB-3xEGFP at the levels of the apical cell surface, centrosomes and nuclei (inverted basket) prior to and following water **C)** or colcemid **D)** injection. The time gap between pre-injection recording and post-injection t = 0 is ∼2’.

## Supplementary Movies

**Supplementary Movie 1**

Live imaging of ventral furrow formation in a representative *Drosophila* embryo (gastrulation stage 6), after water (control, left panel) or Colcemid injection (right panel). Membranes are labeled in green (EGFP-CaaX) and nuclei are labeled in magenta (H2Av-mRFP1). Scale bar is 10 μm.

**Supplementary Movie 2**

Live imaging of cephalic furrow formation along the lateral side of a representative *Drosophila* embryo (gastrulation stage 6), after water (control, left panel) or Colcemid injection (right panel). Membranes are labeled in green (EGFP-CaaX) and nuclei are labeled in magenta (H2Av-mRFP1 [71]). Scale bar is 10 μm.

**Supplementary Movie 3**

Live imaging of dorsal fold formation (mid-sagital view) of a representative *Drosophila* embryo (gastrulation stage 6), after water (control, left panel) or Colcemid injection (right panel). Membranes are labeled in green (EGFP-CaaX) and nuclei are labeled in magenta (H2Av-mRFP1 [71]). Scale bar is μm.

## Supplementary Tables

**Supplementary Table 1**

Summary of the expression pattern of DV fate markers (dpp, sog, snail) in the wild type and DV patterning mutants.

**Supplementary Table 2**

Proteome (LFQ) raw data.

**Supplementary Table 3**

SILAC-Phosphoproteomics raw data.

**Supplementary Table 4**

Linear model implementation: IQR values of the deviation distributions obtained with the systematic exploration of dorsal, lateral and ventral domain proportions.

**Supplementary Table 5**

Log2 deviation values of proteins and phosphosites detected in all genotypes using the dorso-ventral domain proportions: D: 0.4 L: 0.4 V: 0.2. For each experiment (LFQ/Proteomics, SILAC-phosphoproteomics), there is a list of the deviation values of the complete (ANOVA positive and negative) and regulated (ANOVA positive) proteins or phosphosites.

**Supplementary Table 6**

List of proteins and phosphosites within all DV clusters (1-14).

**Supplementary Table 7**

RNA-Proteome comparison: outcome of the RNA-proteome comparison for the list of genes with a mesoderm label in BDGP (https://insitu.fruitfly.org/cgi-bin/ex/insitu.pl), that are also present in the DV clusters.

**Supplementary Table 8**

RNA-Proteome comparison: list of filtered genes from the RNA atlas (155/8924), with their corresponding: DV RNA reference set, proteome DV cluster, Pearson correlation value that allocated each gene to its DV RNA reference set, and the outcome of the RNA-Proteome comparison.

**Supplementary Table 9**

Deviation values of proteins and phosphosites in DV clusters 1-12.

**Supplementary Table 10**

Euclidean distance scores of proteins and phosphosites in DV clusters 1-12.

**Supplementary Table 11**

Proteins and phosphoproteins associated with morphogenesis-related cellular components that are significantly-enriched in diffused networks.

**Supplementary Table 12**

List of proteins and phosphosites with extreme deviations from the linear model.

## Acknowledgements

We thank N.H. Brown, N. Bulgakova, S. Huelsmann, N. Perrimon, V. Riechmann, A. Stathopoulos, D. Stein, S. Roth and A. Wodarz for reagents and fly stocks, N. Lawrence and the Gurdon Institute Imaging Facility for help with 3D-SIM imaging, EMBL Advanced Light Microscopy Facility (ALMF) and CECAD Cologne Imaging Facility for continuous support. Flybase was used throughout the project and is gratefully acknowledged. We also thank Theresa Haunold for 2-photon imaging of myosin in DV patterning mutants, Siegfried Roth for critical discussions, and all members of the Leptin lab for discussions throughout the work.

This work was supported by funding from the European Molecular Biology Organisation (EMBO), the University of Cologne and the Deutsche Forschungsgemeinschaft (grant DFG LE 546/12).

