## Supplementary figures and images for "Differential regulation of the proteome and phosphosproteome along the dorso-ventral axis of the early *Drosophila* embryo"

### Supplementary Figures

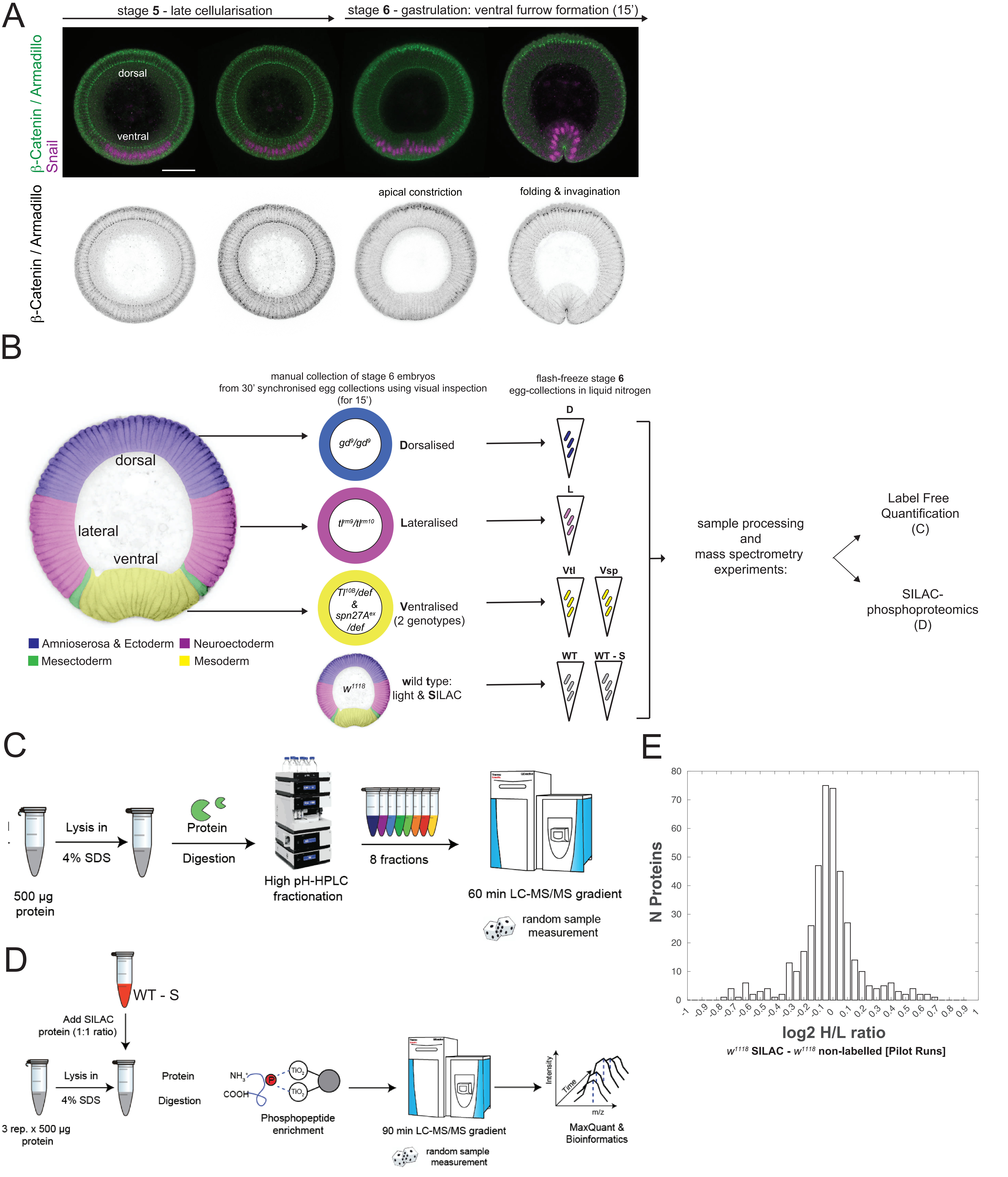

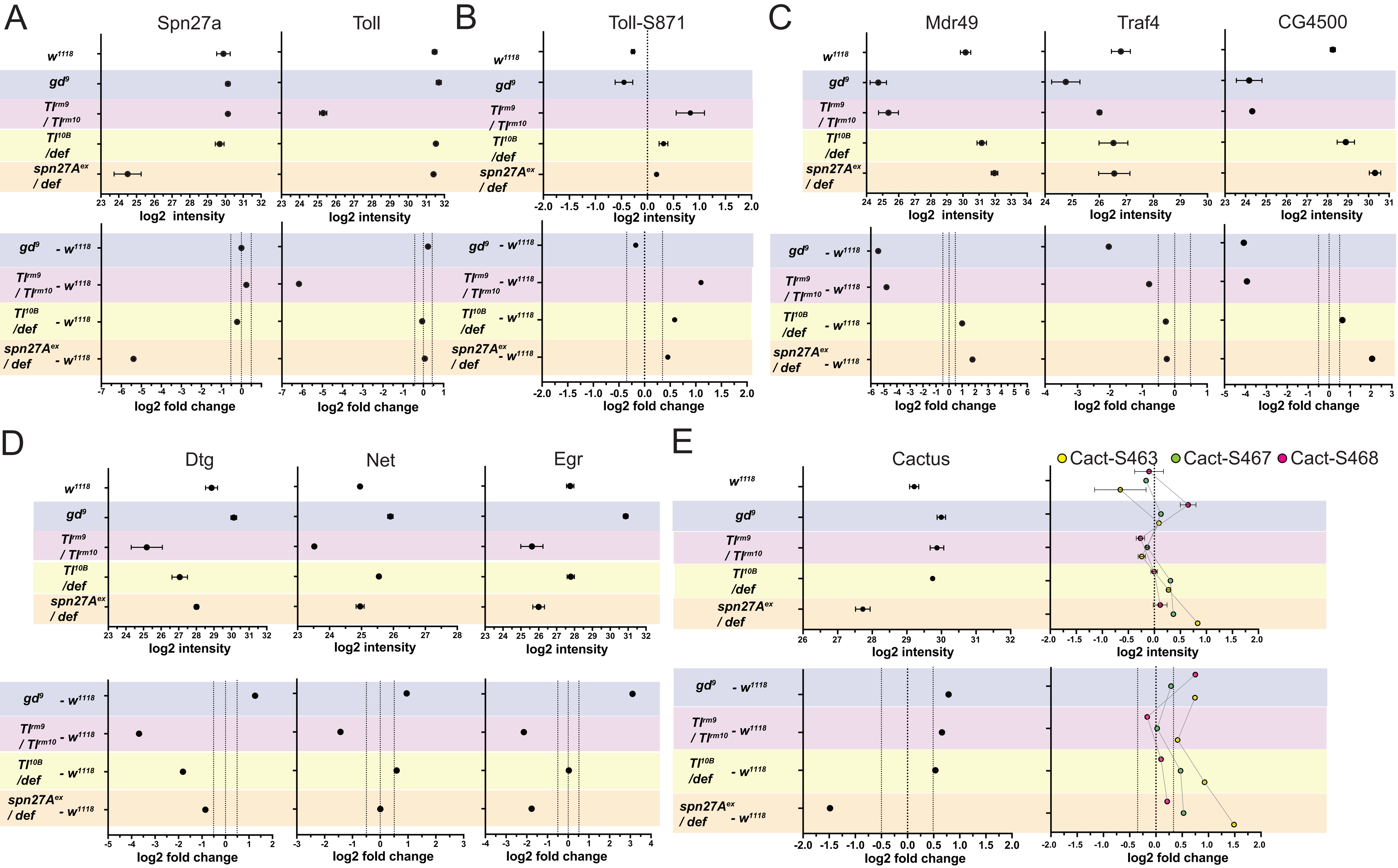

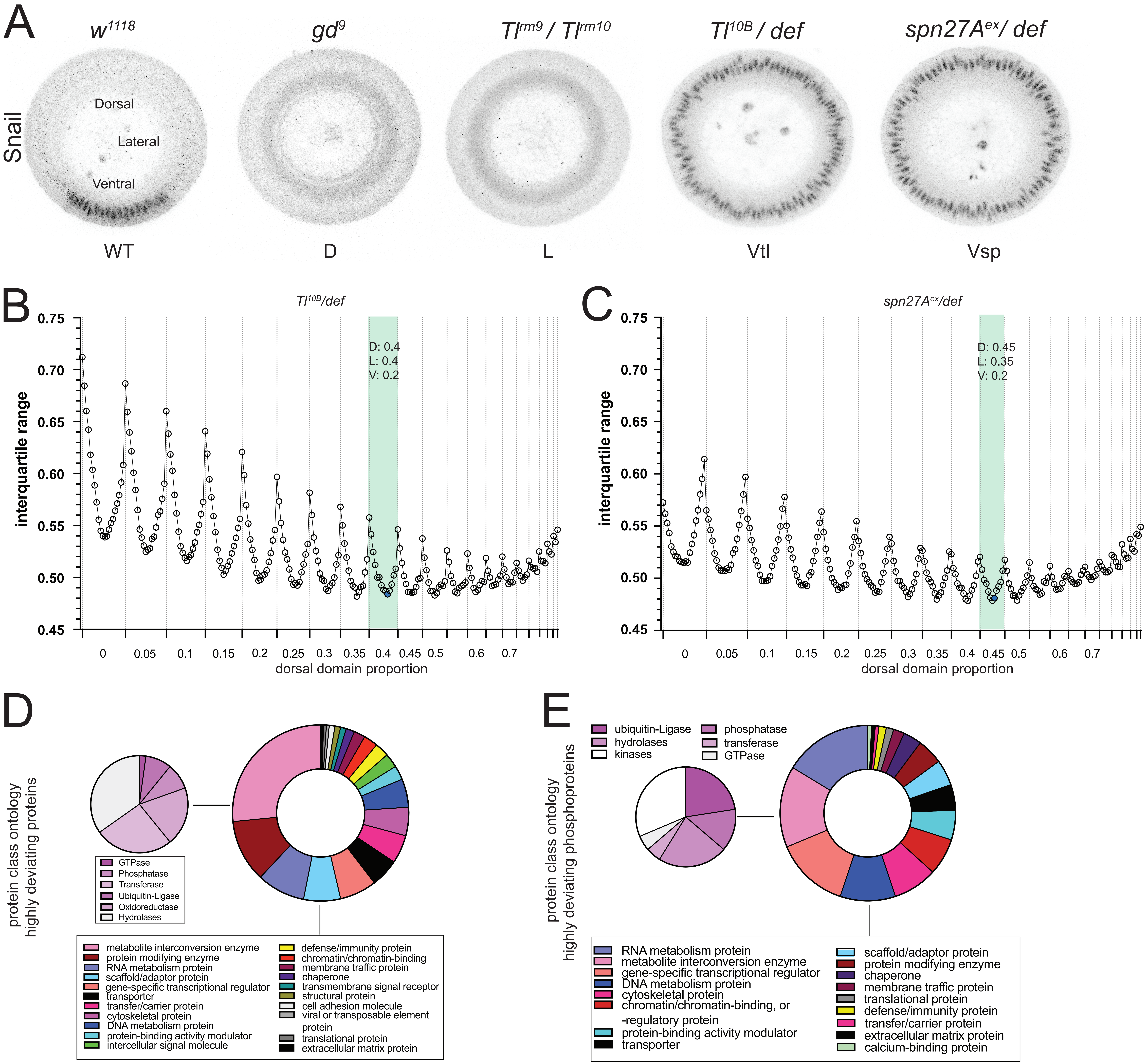

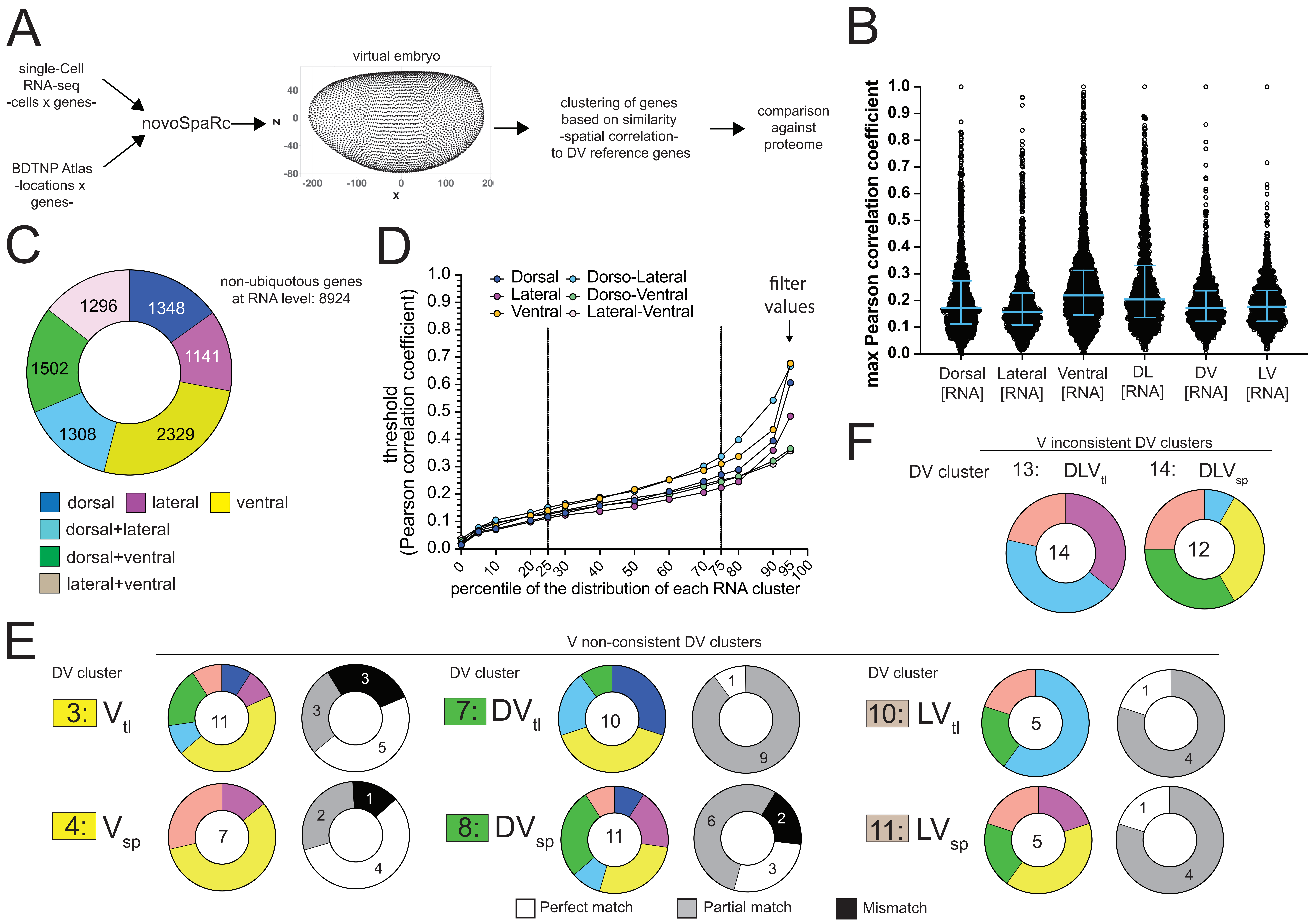

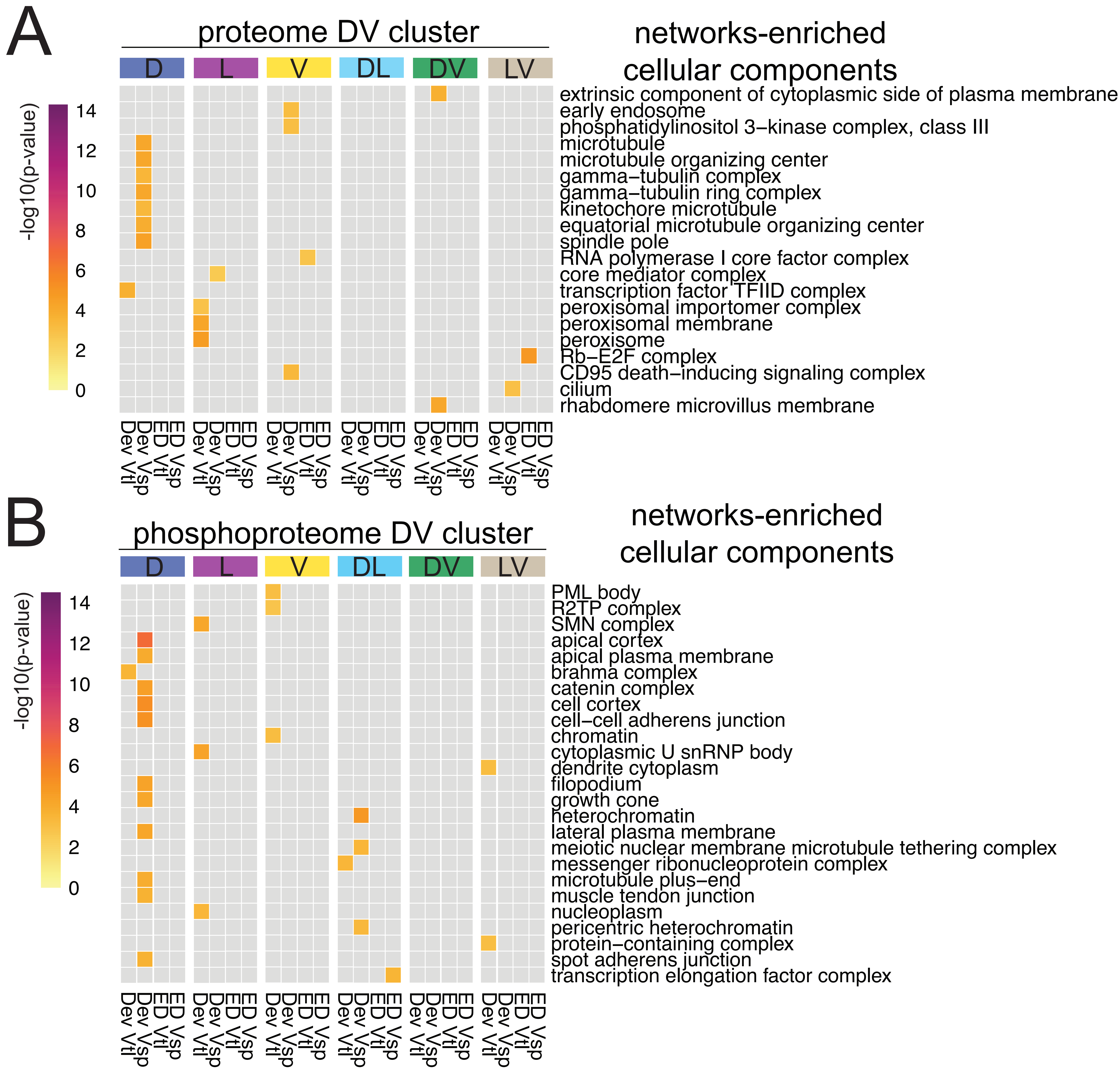

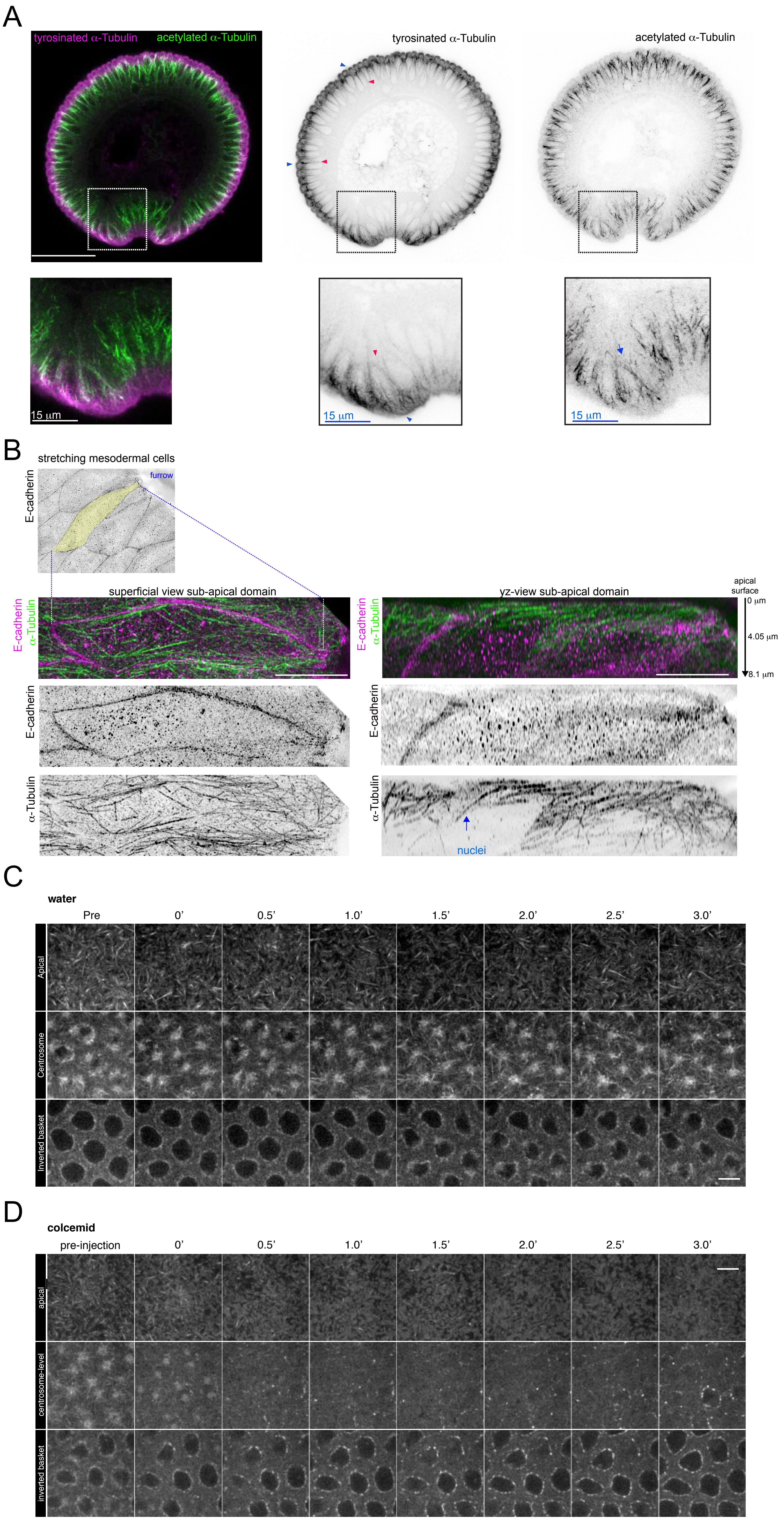
